# Accurate prediction in reconstructed spatial transcriptomes does not ensure valid biological discovery

**DOI:** 10.64898/2026.06.19.733432

**Authors:** Lorenzo Testa, Jing Lei, Kathryn Roeder

## Abstract

Spatial transcriptomics is increasingly extended by computational reconstruction of unmeasured genes from matched single-cell RNA-sequencing references, enabling transcriptome-wide analyses from targeted or sparse assays. Yet downstream analyses typically treat reconstructed expression *as* experimentally observed, overlooking prediction error and latent spatial variation that can misattribute tissue architecture to biological regulation. Here we introduce `TIDEST`, a framework for statistically valid inference on reconstructed spatial transcriptomes. `TIDEST` calibrates reconstructed expression using measured genes and adjusts for latent spatial variation before differential expression analysis. Across realistic spatial tissue simulations, `TIDEST` controls false discoveries where existing approaches fail, while preserving power. Across mouse and human brain, glioblastoma and breast cancer, `TIDEST` changes biological interpretation by correcting misleading differential-expression calls and revealing disease-associated transcriptional programs obscured by spatial confounding. Our results show that prediction accuracy alone is insufficient for reliable biological discovery and establish valid statistical inference as an essential component of reconstructed spatial transcriptomics.

## Introduction

Spatial transcriptomics has transformed the study of tissue biology by measuring gene expression directly within intact tissues, preserving the spatial organization of cells and their local microenvironments [1]. By linking transcriptional programs to tissue architecture, it has become a powerful tool for identifying cellular niches, dissecting disease-associated microenvironments and characterizing developmental processes that cannot be resolved by bulk or dissociated single-cell RNA sequencing [2]. Yet current platforms face an inherent trade-off between spatial resolution and transcriptome coverage. Sequencing-based methods such as 10x Visium [3] and Slide-seq [4] profile thousands of genes but at limited spatial resolution, whereas imaging-based technologies such as 10x Xenium [3], MERFISH [5] and seqFISH [6–8] achieve single-cell or subcellular resolution while measuring only a targeted subset of transcripts. Even transcriptome-wide platforms suffer substantial gene dropout at low sequencing depth, leaving many biologically relevant genes incompletely observed. These limitations complicate one of the central goals of spatial studies: identifying genes whose expression differs across tissue regions, cellular neighborhoods and disease states.

To overcome incomplete coverage, a growing class of deep learning methods reconstructs unmeasured or sparsely observed expression by integrating spatial data with matched single-cell RNA-sequencing references [9–22]. Approaches such as `Tangram` [10] and `gimVI` [22] exploit shared transcriptional structure to infer transcriptome-wide spatial expression, substantially expanding the scope of downstream analyses, and have become widely used both for interpreting targeted imaging assays and for recovering expression lost to technical dropout. Reconstructed values, however, are predictions rather than direct molecular measurements. Current differential expression analyses typically either discard reconstructed genes or analyze predicted expression as though it were observed, overlooking the uncertainty introduced during reconstruction [23]. Compounding this, neighboring locations often share transcriptional programs driven by tissue architecture, cell-type composition and microenvironmental gradients, so apparent differences between biological groups may reflect where cells sit rather than the condition of interest. Together, prediction uncertainty and this latent spatial structure can produce misleading expression differences and erode confidence in the resulting biology. As a result, no existing method supports reliable differential expression analysis on reconstructed spatial transcriptomes.

Here we introduce *Testing for Imputed Differential Expression in Spatial Transcriptomics* (`TIDEST`), a framework that enables reliable differential expression analysis on reconstructed spatial transcriptomes. Unlike existing methods, which either discard reconstructed genes or treat predicted expression as if it were directly observed, `TIDEST` first calibrates reconstructed expression using measured transcripts to reduce systematic prediction error. It then estimates and adjusts for latent spatial variation arising from tissue architecture and cellular organization before testing for differential expression, providing a statistically principled foundation for inference. In realistic simulations of spatial tissue architecture, widely used methods including `SpaGCN` [24], `DESpace` [25] and `SpatialGEE` [26] fail to maintain appropriate false-positive rates when reconstruction error or spatial structure is present, whereas `TIDEST` achieves reliable error control while preserving power. Applied to mouse and human brain, glioblastoma and breast cancer, `TIDEST` corrects misleading differential-expression calls and reveals spatial genetic and disease-associated transcriptional programs across diverse biological settings – including cortical lamination, genetic risk for neuropsychiatric traits, glioblastoma cell states, and breast cancer invasion – that are missed or mischaracterized by conventional analyses.

## Results

### Overview of TIDEST

`TIDEST` is a framework for transcriptome-wide differential expression testing in spatial transcriptomics datasets where some genes are reconstructed from a scRNA-seq reference. The workflow takes as input a ST dataset, a matched scRNA-seq reference, and the predicted spatial expression generated by an imputation model (Fig. **1a**). `TIDEST` then refines the reconstructed expression profiles using information from measured genes (Fig. **1b**), identifies latent spatial variation that may confound biological comparisons (Fig. **1c**), and performs DE testing between predefined biological conditions (Fig. **1d**). The output is a set of effect estimates, confidence intervals, and significance measures for both observed and reconstructed genes, enabling transcriptome-wide DE analysis while accounting for uncertainty introduced during spatial reconstruction.

**Figure 1:**
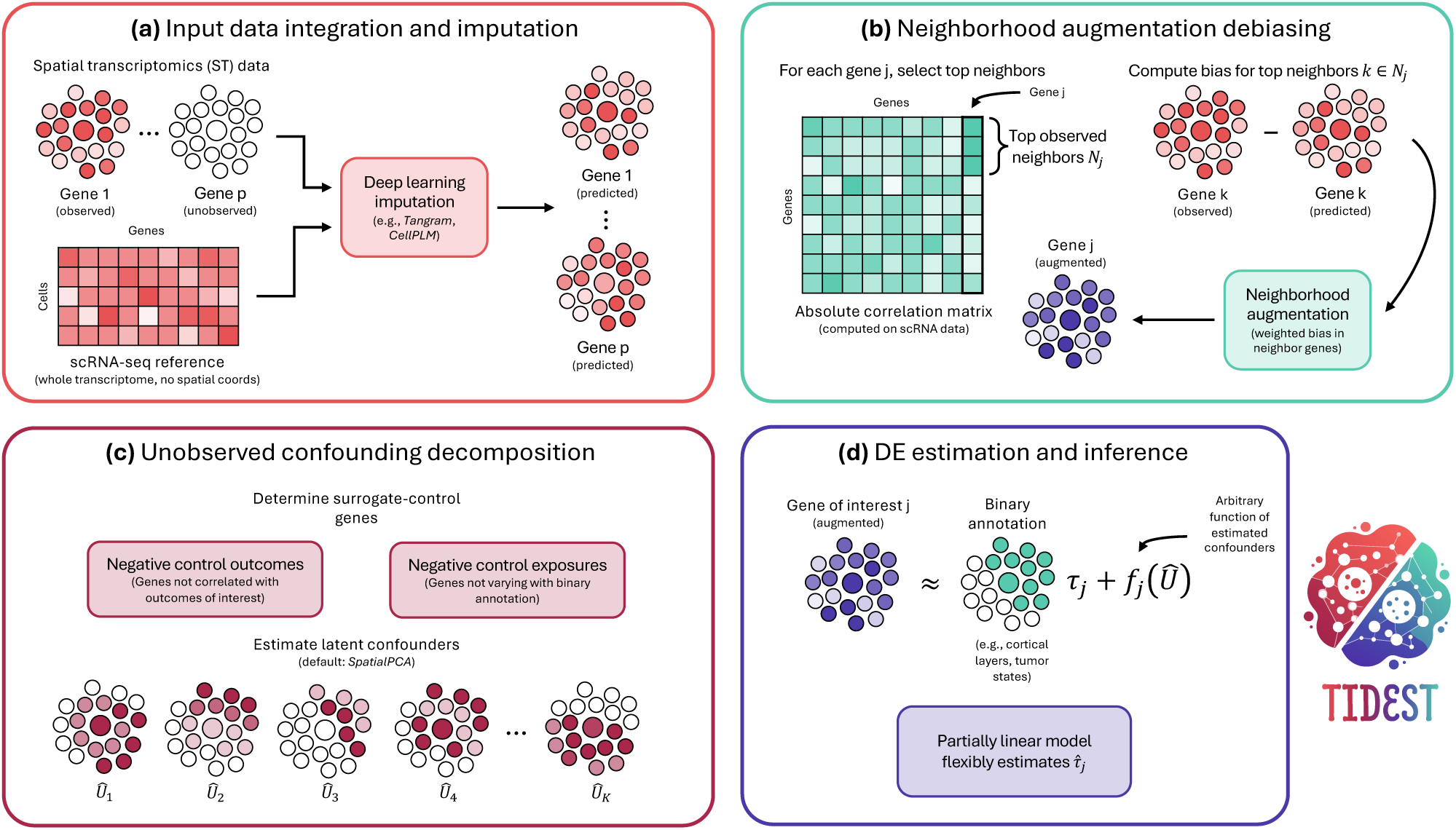
Overview of TIDEST. **(a)** `TIDEST` takes as input a spatial transcriptomics (ST) dataset, a matched scRNA-seq reference and the reconstructed transcriptome generated by an imputation model. The scRNA-seq reference provides transcriptome-wide information, whereas the ST data provide spatial localization. **(b)** `TIDEST` refines the reconstructed expression profiles using information from measured genes. For each target gene, the method identifies highly correlated observed genes and uses their prediction errors to correct systematic biases in the reconstructed expression. This produces an augmented expression profile that is less sensitive to errors introduced during imputation. **(c)** `TIDEST` identifies large-scale spatial patterns that may influence gene expression independently of the biological comparison of interest. Examples include anatomical structure, cell-type composition, microenvironmental gradients, and technical effects. These latent patterns are estimated from a set of control genes and summarized as a low-dimensional representation of spatial variation. **(d)** `TIDEST` performs differential expression testing between predefined biological groups while accounting for the estimated spatial variation. The output consists of effect estimates, confidence intervals, and significance measures for both observed and reconstructed genes, enabling transcriptome-wide DE analysis from spatially reconstructed transcriptomes.

The first stage addresses prediction uncertainty via a *debiasing* procedure (Fig. **1b**) [27, 28]. Common practice treats imputed values *Ỹ_ij_* for spatial coordinate *i* and gene *j* as true observations, ignoring the error introduced during reconstruction. `TIDEST` instead uses the observed residuals of highly correlated measured genes to construct an *augmented* outcome *Ŷ_ij_* for each target gene (see Methods), reducing systematic prediction errors and improving robustness to biases introduced by the underlying imputation model.

The second stage addresses unobserved confounding by estimating latent sources of variation from surrogate-control outcomes (Fig. **1c**). In spatial transcriptomics, factors other than the biological comparison of interest can create apparent expression differences between groups. For example, anatomical structure, cell-type composition, microenvironmental gradients or technical effects may vary across a tissue and influence gene expression independently of the condition being studied [29]. `TIDEST` estimates these latent patterns (by default using `SpatialPCA` [30]) and incorporates them into downstream analyses, reducing the risk that such factors are mistaken for true differential expression signals.

The final stage performs DE testing while accounting for the estimated latent variation (Fig. **1d**). `TIDEST` fits a partially linear model [31] relating the augmented outcomes to the biological condition of interest while flexibly adjusting for the estimated spatial confounders. Because inference is performed on the augmented outcomes rather than the raw predictions, `TIDEST` simultaneously addresses prediction uncertainty and unobserved spatial variation, enabling more reliable identification of differentially expressed genes across biological conditions.

### Simulation study provides strong support for TIDEST

We validated `TIDEST` on fully synthetic data in which ground-truth treatment effects, spatial confounders, and imputation errors are known by design (see Methods). An example scRNA-seq reference, spatial treatment *A*, unobserved spatial confounder *U*, and observed gene expression *Y* are shown in Fig. **2a-d**. We compared `TIDEST` against a two-sample *t*-test and three representative methods for differential expression analysis in spatial transcriptomics [23]: `SpaGCN` [24] (Wilcoxon rank-sum test, also used in `Seurat` [32]), `DESpace` [25] (negative binomial generalized linear model), and `SpatialGEE` [26] (generalized estimating equations).

**Figure 2:**
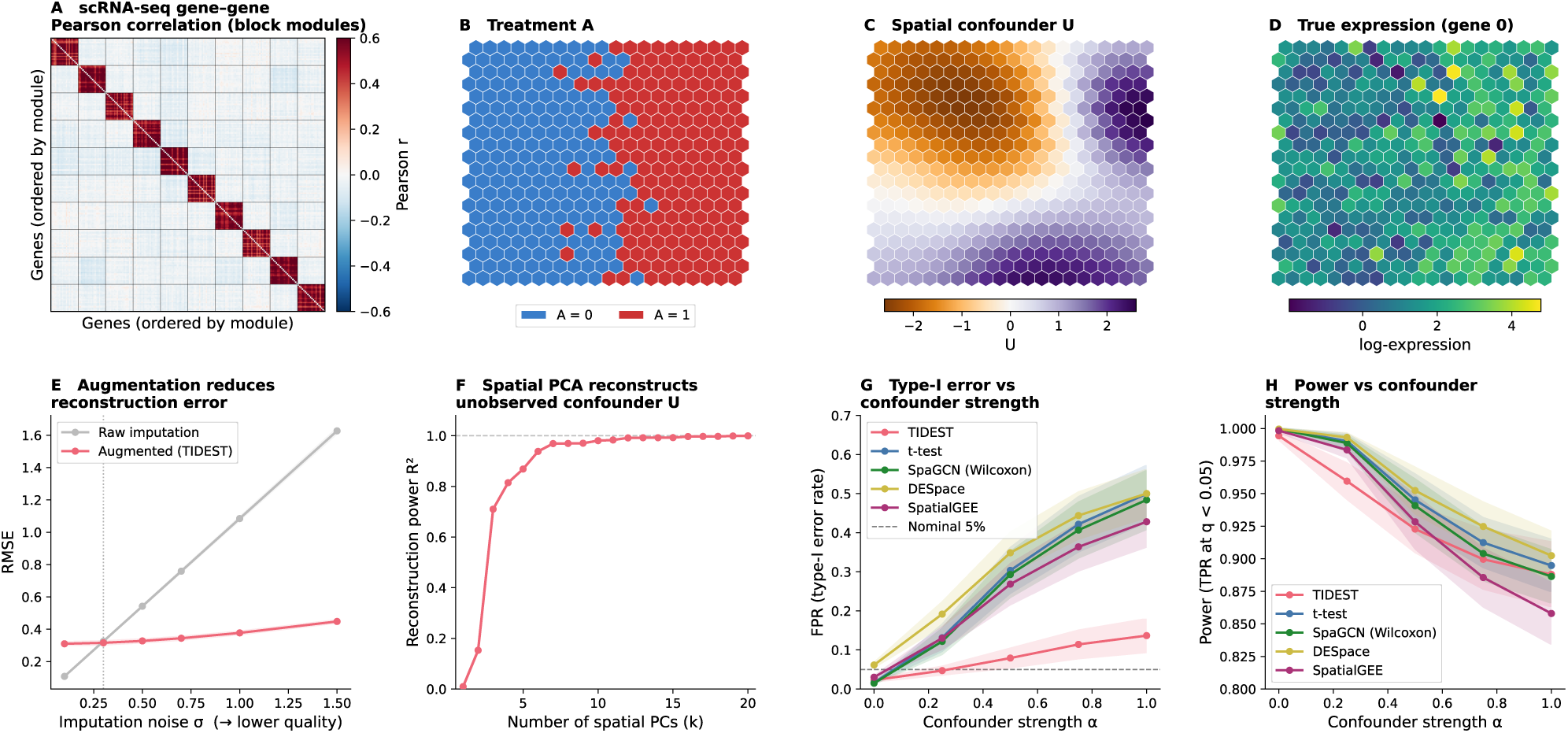
Simulation study validating TIDEST. **(a)** Block-diagonal Pearson correlation matrix of the synthetic scRNA-seq reference, computed from 2000 cells across 10 co-expression modules (200 genes). The block structure is the informative gene-gene correlation exploited by `TIDEST` augmentation strategy. **(b)** Binary treatment assignment *A* (two-region DGP, *n* = 400 spots on a regular hexagonal grid). **(c)** Unobserved spatial confounder *U*, drawn as a Gaussian random field with length-scale A = 0.3. In this simulation setting, *U* is spatially smooth and independent of *A*, but its interaction with gene loadings *β_j_* induces systematic differences between treatment regions that naive comparisons absorb into the estimated treatment effect. **(d)** True log-expression of a representative DE gene, combining the treatment effect *τ_j_ A_i_* and the confounder contribution *αβ_j_ U_i_*. **(e)** Reconstruction quality of the augmented pseudo-outcome. RMSE of the raw imputed prediction (*Ỹ*, gray) and the augmented outcome (*Ŷ*, red) against the true latent expression *Y*, as a function of imputation noise σ_imp_ (lower σ_imp_ means better imputer). The dashed vertical line marks the crossover (σ_imp_ ≈ 0.3) below which raw imputation is sufficient; above this threshold the augmented RMSE is bounded between 0.31 and 0.45 regardless of imputer quality. **(f)** Confounder reconstruction power. Coefficient of determination *R*^2^ of *U* as a function of the number of spatial PCA eigenvectors *k* retained in the confounder matrix *U*, computed on the illustration replicate. Three eigenvectors capture 71% of the confounder variance; 10 eigenvectors reach 98%. **(g)** Type-I error rate (FPR at BH *q <* 0.05) vs α (50 replicates per α level). The dashed line marks the nominal 5% level. All competing methods inflate FPR steeply with confounding; `TIDEST` maintains substantially lower FPR across all settings. Shaded bands are 95% confidence intervals across replicates. **(h)** Power (TPR at BH *q <* 0.05) vs confounder strength α (50 replicates per α level).

Augmentation provides no benefit when prediction is already highly accurate, but substantially improves reconstruction throughout the error range typical of current prediction methods (Fig. **2e**). Importantly, augmentation remains stable across a wide range of imputation quality, indicating that downstream inference need not depend critically on the accuracy of the prediction model. The learned confounder representation *Û* approximates the unobserved field *U* well (Fig. **2f**) – even three eigenvectors capture most confounder variation, and increasing *k* yields near-complete removal of spatial confounding.

Existing methods rapidly lose false-discovery control as spatial confounding increases, whereas `TIDEST` remains substantially more robust (Fig. **2g**). At α = 1, all competing methods exhibit approximately ten-fold inflation in false discoveries (43–50% FPR), whereas `TIDEST` limits FPR to 13.7%. We next examine sensitivity (TPR at BH *q <* 0.05): at α = 0 all methods achieve near-perfect power, and at α = 1.0 power converges toward 85-90% across methods (Fig. **2h**). However, the apparent advantage of the *t*-test and `DESpace` [25] over `TIDEST` at high α is an artifact of inflated FPR – the BH threshold admits more false positives while true positives barely increase. This is evident from the AUC (Supp. Fig. B.1), which evaluates how well each method separates truly DE genes from null genes across all possible significance thresholds. At α = 0 all methods achieve AUC 1, while at α = 1.0 `TIDEST` achieves a 10 percentage-point advantage.

Together, these simulations demonstrate that accurate reconstruction does not guarantee valid biological inference. Rather, prediction uncertainty interacting with latent spatial variation produces systematic false discoveries unless explicitly modeled. By calibrating reconstructed expression and adjusting for latent spatial variation, `TIDEST` restores reliable differential-expression inference across a wide range of imputation quality.

### Correcting reconstruction bias resolves misleading laminar differential expression in the mouse neocortex

We first evaluated whether `TIDEST` could improve the biological interpretation of reconstructed spatial transcriptomes in the adult mouse neocortex. We analyzed a Visium spatial transcriptomics section [33] comprising 297 annotated spots from superficial (L2/3/4; *n* = 126) and deep (L5/6; *n* = 171) cortical layers (Fig. **3a**). To mitigate Visium dropout, we projected a matched scRNA-seq reference (21697 cells) onto the tissue using `Tangram` [10, 34]. Because cortical gene expression varies continuously across depth [35, 36], `TIDEST` accounted for this spatial structure when testing differential expression between superficial and deep layers (see Methods).

**Figure 3:**
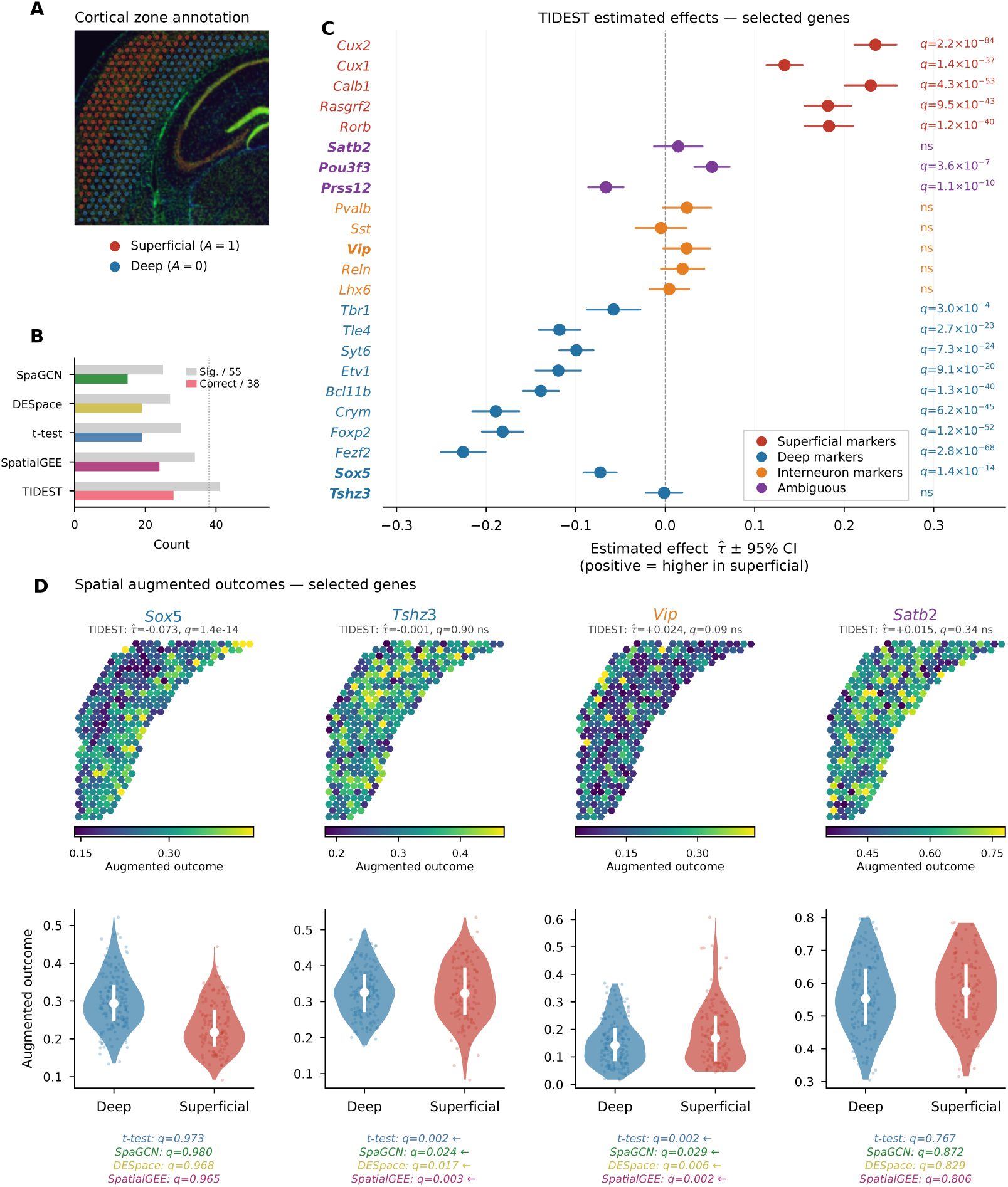
Mouse brain application. **(a)** Visium section of the adult mouse neocortex with cortical zone annotation overlaid as colored hexagonal spots (red: superficial layers L2/3/4, *A* = 1; blue: deep layers L5/6, *A* = 0; 297 spots total). **(b)** Method comparison summary. Grouped bars show, for each method, the number of genes called significant out of all 55 (gray) and the number that are both significant and correctly directioned out of the 38 genes with an established expected direction (colored). `TIDEST` achieves the highest correct count (28/38). **(c)** `TIDEST` estimated effects (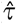 *±* 95% CI) for 23 selected genes. Color indicates gene category: red = superficial markers, blue = deep markers, orange = interneuron markers (negative controls), purple = ambiguous annotation. BH-adjusted *q*-values are shown to the right of each estimate. **(d, top row)** Spatial augmented outcome maps for four key genes (*Sox5*, *Tshz3*, *Vip*, *Satb2*). The subtitle reports the `TIDEST` estimate and BH-adjusted *q*-value. **(d, bottom row)** Deep vs. superficial augmented outcome distributions for the same four genes. Violins show the full distribution; the white bar marks the interquartile range; the white dot marks the median. BH-adjusted *q*-values for all four competing methods (`SpaGCN`, `DESpace`, *t*-test, `SpatialGEE`) are annotated below each violin. *Sox5*: `TIDEST` detects a deep signal (*q* = 1.4 × 10^−14^) that all competitors miss (*q* ≥ 0.97). *Tshz3*: `TIDEST` returns a null result (*q* = 0.90) while all competitors produce significant estimates in the wrong biological direction (*q* = 0.024). *Vip*: interneuron marker that may not differ by layer; `TIDEST` is non-significant (*q* = 0.09) while all competitors produce significant results (*q* = 0.029). *Satb2*: expressed across both layers; all methods agree on non-significance.

Several genes illustrate how accounting for reconstruction bias and latent spatial variation changes biological interpretation (Fig. **3d**). *Sox5*, a canonical deep-layer marker [37, 38], is detected by `TIDEST` (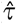 = −0.073, *q* = 1.4 × 10^−14^) but missed by all four competitors (*q* ≥ 0.965). Conversely, *Tshz3*, a deep-layer gene [39], is returned as non-significant by `TIDEST` (*q* = 0.903), while all four competing methods instead identified significant differential expression in the opposite biological direction: *t*-test calls it superficial (*q* = 0.002), as do `SpatialGEE` (*q* = 0.003), `DESpace` (*q* = 0.017), and `SpaGCN` (*q* = 0.024). The clearest specificity advantage concerns interneuron markers, which are distributed across cortical layers and may not systematically differ between zones. *Vip*, a canonical interneuron marker [40], is called significant by every competitor (*t*-test *q* = 0.002, `SpatialGEE` *q* = 0.002, `DESpace` *q* = 0.006, `SpaGCN` *q* = 0.029), but `TIDEST` returns *q* = 0.090 (non-significant). These examples illustrate that statistically valid inference changes biological interpretation rather than simply increasing statistical power.

To evaluate this improvement systematically, we assembled a curated panel of 55 cortical genes comprising 38 established layer markers with known directionality, five interneuron markers expected to show no laminar bias, and 12 genes with uncertain layer preference. Overall, `TIDEST` identified 41/55 (74.5%) as significantly differentially expressed (BH *q <* 0.05) (Fig. **3b**). The strongest superficial signal was *Cux2* (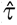 = +0.235, *q* = 2.2 10^−84^), a well-established upper-layer marker [41]; the strongest deep signal was *Fezf2* (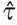 = 0.226, *q* = 2.8 10^−68^), a canonical regulator of corticospinal neuron identity [42, 43] (Fig. **3c**; further examples in Supp. Material). Compared with four competitors on the same gene panel (Fig. **3b**; Supp. Fig. C.3), `TIDEST` recovered the most correct-direction significant genes (28 of 38), versus `SpatialGEE` (24), *t*-test (19), `DESpace` (19), and `SpaGCN` (15). The improvement was most apparent for genes with moderate effect sizes or gradual laminar gradients, where explicitly modeling spatial dependence reduced both false negatives and biologically misleading discoveries.

`TIDEST` also recovered signals consistent with recent transcriptomic evidence that refines earlier functional annotations (Fig. **3c**); two further examples (*Pou3f3*, *Prss12*) are detailed in the Supp. Material. *Satb2* was originally characterized as a superficial marker required for callosal projection neuron identity in L2/3 [44, 45], although its immunoreactivity has also been observed in L6, challenging a strictly superficial interpretation [46, 47]. `TIDEST` returns a non-significant estimate (*q* = 0.339; Fig. **3d**), consistent with expression across both zones – a null result shared by all competing methods. Likewise, the remaining four interneuron markers were non-significant in `TIDEST` (Supp. Table C.1), suggesting that apparent layer associations detected by competing methods may instead reflect local interneuron clustering rather than robust laminar enrichment.

### Genetic risk for neuropsychiatric and cognitive traits converges on Layer 2/3 of the human DLPFC

We applied `TIDEST` to 12 Visium sections of the human dorsolateral prefrontal cortex (DLPFC) from three neurotypical donors [48], asking whether genes nominated by GWAS Locus-to-Gene (L2G) analyses for three brain-related traits – educational attainment (EA) [49], schizophrenia (SCZ) [50], and intelligence (IQ) [51] – show layer-specific spatial enrichment relative to expression-matched background genes (Fig. **4a,b**). We tested each of five manually annotated cortical layers (Layer 1, Layer 2/3, Layer 4, Layer 5, Layer 6) against all remaining retained layers and combined section-specific estimates across the 12 sections using random-effects meta-analysis (see Methods).

**Figure 4:**
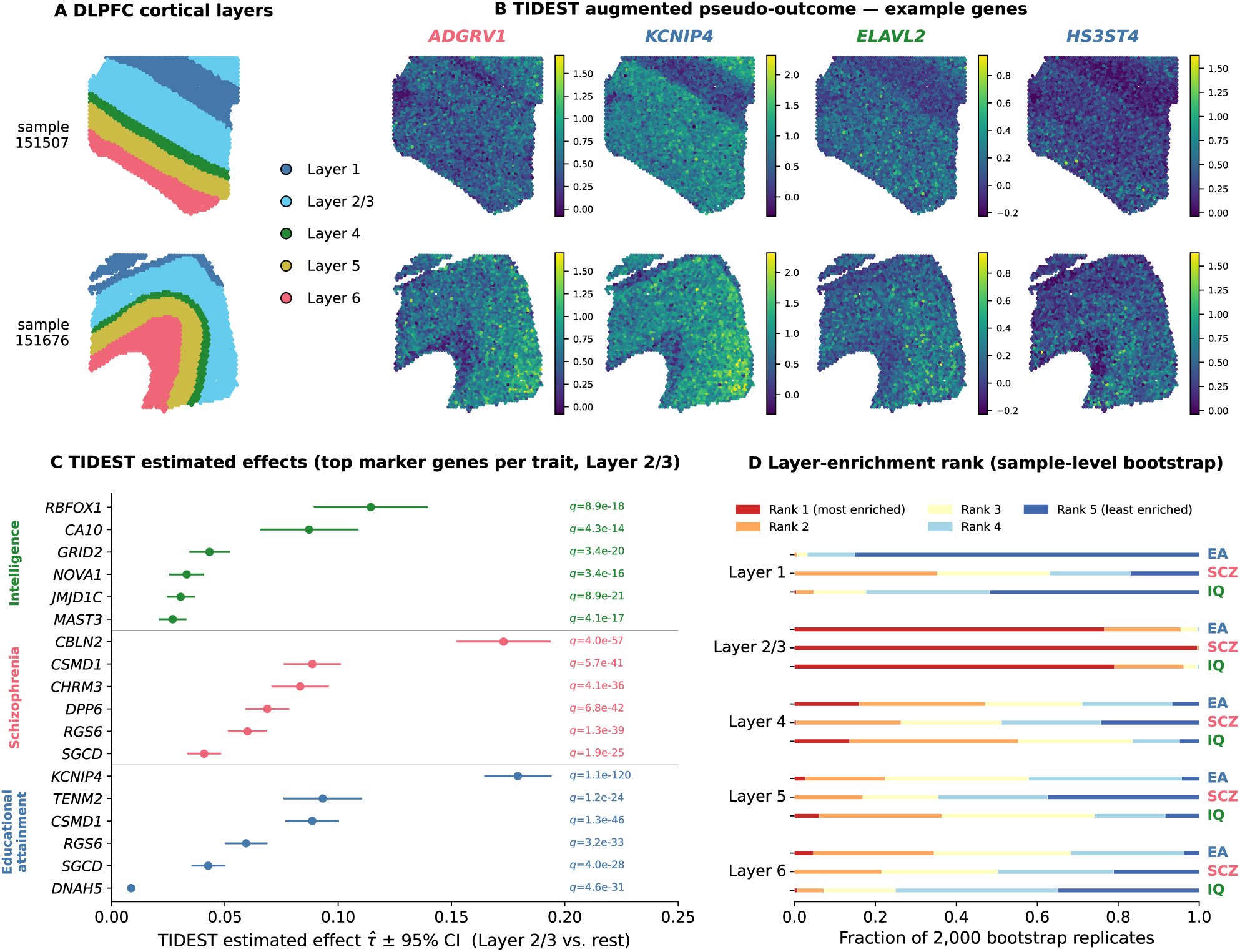
Human brain application. **(a, b)** Manual cortical-layer annotation (**a**) and `TIDEST` augmented spatial expression maps for four representative marker genes (**b**; *ADGRV1*, *KCNIP4*, *ELAVL2*, *HS3ST4*) across two representative DLPFC sections (151507, 151676), illustrating that GWAS-nominated marker genes for different traits peak in different cortical layers. **(c)** `TIDEST` estimated effects (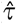 *±* 95% CI, Layer 2/3 vs. all other retained layers) for the top 6 marker genes (by pooled *q*-value) for each of three GWAS traits – educational attainment (EA), schizophrenia (SCZ), and intelligence (IQ) – fitted independently in each of the 12 DLPFC sections and combined by DerSimonian-Laird random-effects meta-analysis. *q*-values (BH-adjusted within Layer 2/3) are shown to the right of each estimate. **(d)** Stability of the layer ranking by trait-gene enrichment. For each trait and cortical layer, we resampled the 12 sections with replacement (2000 bootstrap replicates), recomputed the random-effects meta-analysis, and ranked the 5 layers by the common language effect size comparing trait genes to matched background genes (restricted to genes with positive 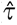). Stacked bars show, for each layer and trait, the fraction of the 2000 replicates in which that layer was assigned each rank (rank 1 = most enriched, rank 5 = least enriched). Layer 2/3 is the most consistently enriched layer for all three traits.

A strikingly consistent pattern emerged across all three traits: Layer 2/3 showed the strongest enrichment of GWAS-nominated genes. More trait genes reached genome-wide-adjusted significance (BH *q <* 0.05) in Layer 2/3 than in any other layer, comprising 117/199 (58.8%) genes for EA, 141/221 (63.8%) for SCZ, and 78/146 (53.4%) for IQ (Fig. **4c**). This convergence was observed despite the three trait panels being derived from non-overlapping discovery GWAS with distinct, though correlated, genetic architectures and independently constructed expression-matched background panels.

To determine whether Layer 2/3 enrichment represented a robust cross-section pattern rather than being driven by a small number of influential tissue sections, we performed a sample-level bootstrap. We resampled the 12 sections with replacement, recomputed the random-effects meta-analysis, and ranked the five layers by the common-language effect size comparing trait genes with expression-matched background genes (restricted to positive 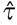 values) across 2000 replicates. Layer 2/3 was ranked as the most enriched layer in 76.6% of replicates for EA, 99.6% for SCZ, and 79.0% for IQ (Fig. **4d**), making it the modal top-ranked layer by a wide margin for every trait. Conversely, Layer 1 was consistently identified as the least-enriched layer for EA and IQ (85.2% and 51.8% of replicates ranked it last, respectively), whereas for SCZ the least-enriched rank was more evenly distributed between Layers 4 and 5. Together, these results identify Layer 2/3 as a reproducible point of convergence for genetic signals associated with cognitive and neuropsychiatric traits in the human DLPFC.

The strongest Layer 2/3-enriched gene differed across phenotypes, suggesting that this cross-trait convergence encompasses distinct molecular programs. For EA, the strongest signal was *KCNIP4* (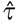 = 0.179, *q* = 1.1 10^−120^), which encodes a calcium-binding subunit of native Kv4 potassium channel complexes involved in neuronal excitability [52, 53]. For SCZ, *CBLN2* (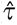 = 0.173, *q* = 4.0 10^−57^) encodes a synaptic organizer implicated in schizophrenia and related neuropsychiatric disorders [54, 55]. For IQ, *RBFOX1* (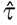 = 0.114, *q* = 8.9 10^−18^) encodes a neuronal splicing factor regulating transcripts involved in neuronal excitability and has been linked to learning and memory [56]. Thus, independent genetic architectures converge spatially on Layer 2/3 while implicating distinct genes involved in neuronal excitability, synaptic organization, and RNA regulation.

### Spatial confounding obscures tumor-intrinsic transcriptional programs at the invasive margin of human glioblastoma

IDH-wildtype glioblastoma (GBM) is a genetically defined, highly heterogeneous primary brain tumor in which malignant cells coexist with and infiltrate normal brain tissue, complicating the interpretation of spatial expression differences. The cellular tumor (CT) and leading edge (LE) provide a particularly stringent setting for spatial differential expression: CT regions are densely packed with neoplastic cells, whereas LE regions contain infiltrating tumor cells interdigitated with resident brain tissue. Consequently, expression differences between these compartments can reflect both tumor-intrinsic transcriptional programs and changes in local cellular composition, making it difficult to distinguish malignant programs from signals contributed by the surrounding brain microenvironment.

We therefore analyzed 26 10x Visium sections spanning three independent patient cohorts [57]. We fitted `TIDEST` independently to each section using the Ivy Glioblastoma Atlas Project (IvyGAP) annotation [58], comparing spots assigned to cellular tumor (CT) with those at the leading edge (LE), with positive 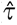 values indicating higher expression at the invasion front (Fig. **5a**). To reduce the impact of dropout, expression was imputed using `Tangram` [10] with an external scRNA-seq reference from a disjoint cohort [59]. We assembled a curated panel of 90 genes (Supp. Table D.2) spanning GBM cell states, neuronal and myelin markers, invasion mediators, immune genes, and metabolic programs, and combined section-specific estimates using random-effects meta-analysis.

**Figure 5:**
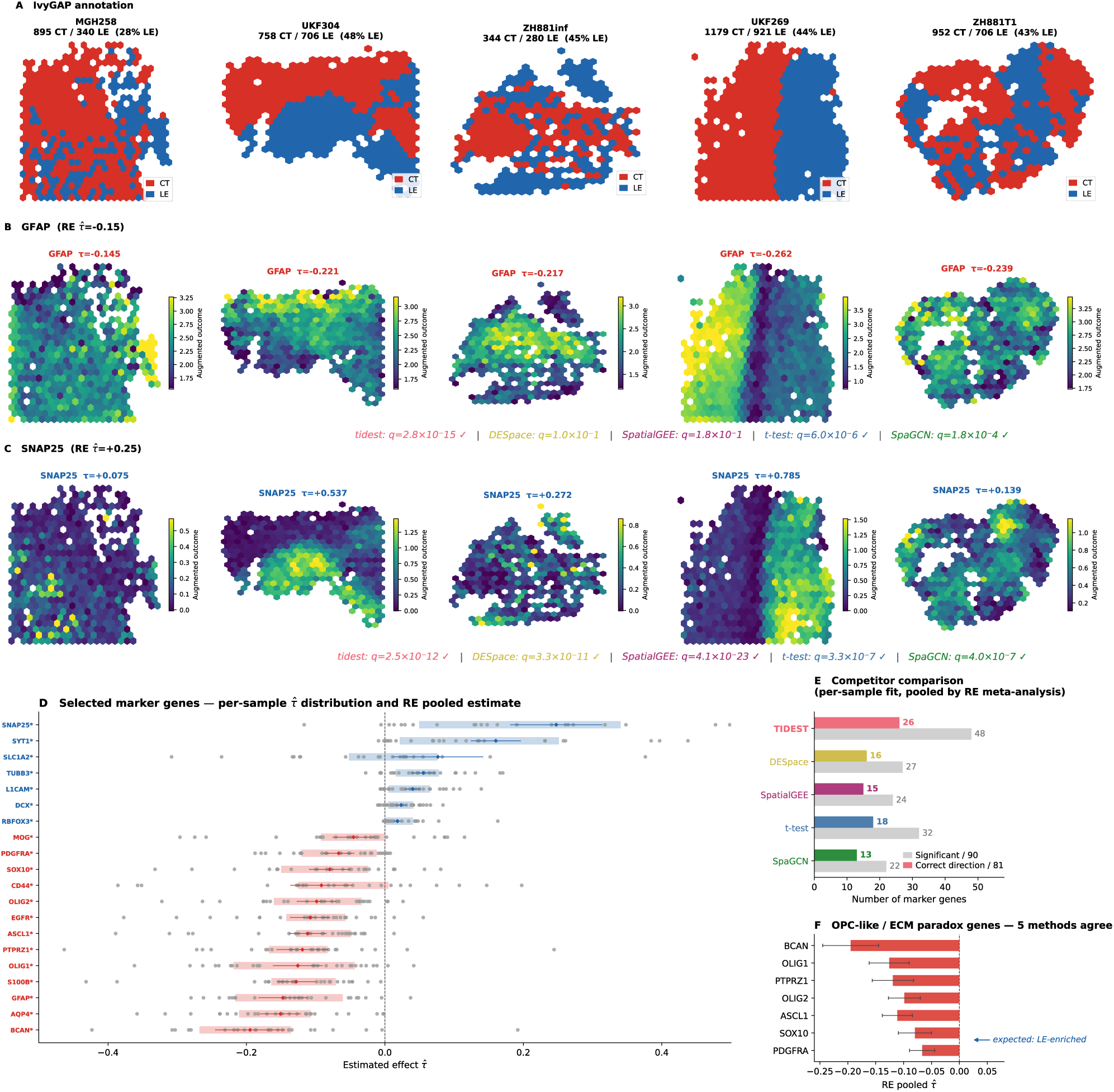
Human glioblastoma application. **(a–c)** IvyGAP histological annotation (**a**; CT in red, LE in blue) and augmented spatial expression maps of *GFAP* (**b**) and *SNAP25* (**c**) across five representative GBM sections (MGH258, UKF304, ZH881inf, UKF269, ZH881T1), illustrating the spatial cell-composition gradient that `TIDEST` disentangles from genuine tumor biology. Per-sample 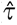 values are shown above each map; pooled random-effects estimates are given in the row titles. Colored annotations below the maps report each method’s *q*-value for that gene, illustrating `TIDEST`’s power advantage on representative markers. **(d)** Per-sample 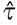 estimates (dots) and random-effects pooled estimates with 95% confidence intervals (diamonds and shaded bands) for a curated panel of marker genes, ordered by pooled 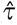 from strongly cellular-tumor (CT)-enriched (left, red) to strongly leading-edge (LE)-enriched (right, blue). **(e)** Number of the 90 panel genes called significant (BH *q <* 0.05; gray bars) and, of the 81 genes with an *a priori* expected CT/LE direction, the number called significant with the correct sign (colored bars), for `TIDEST` and four competing methods (*t*-test, `SpaGCN`, `DESpace`, `SpatialGEE`), each fitted independently per sample across all 26 sections and pooled with the same DerSimonian-Laird random-effects meta-analysis. **(f)** Random-effects pooled 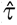 (with 95% confidence intervals) for the seven OPC-like and ECM “paradox” genes (*BCAN*, *OLIG1*, *PTPRZ1*, *OLIG2*, *ASCL1*, *SOX10*, *PDGFRA*). All seven are robustly CT-enriched (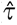 *<* 0, BH *q <* 10^−6^ for every gene) and all five tested methods agree unanimously on the CT-enriched sign for every gene.

`TIDEST` resolved a clear transcriptional contrast between the tumor core and invasive margin. Among established GBM driver genes, *EGFR* – amplified in approximately 40% of IDH-wildtype tumors – showed robust CT enrichment (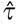 = 0.108, *q* = 2.3 10^−12^, significant in 73% of samples), demonstrating recovery of the expected spatial localization of a canonical genetic driver despite extensive cellular heterogeneity. *CD44*, one of the best-established mesenchymal GBM markers, was similarly CT-enriched (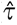 = 0.092, *q* = 1.4 10^−4^, 77%) [60]. The strongest CT-enriched signals corresponded to genes characteristic of the astrocyte-like transcriptional state identified in single-cell analyses of genetically diverse GBMs [61], including *AQP4* (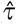 = 0.151, *q* = 2.4 10^−22^, 85%), *S100B* (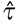 = 0.128, *q* = 4.0 10^−18^, 92%), and *GFAP* (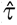 = 0.147, *q* = 2.8 10^−15^, 73%), consistent with the high abundance of astrocyte-like malignant cells in the tumor core [62] (Fig. **5b**).

In contrast, the strongest LE-enriched signals reflected the distinct biology of the invasive margin. The synaptic vesicle genes *SYT1* and *SNAP25* were among the most prominent positive-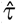 signals (*SYT1*: 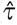 = +0.160, *q* = 5.7 10^−17^, 73%; *SNAP25*: 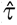 = +0.247, *q* = 2.5 10^−12^, 77%; Fig. **5c**), and several additional neuronal and astrocytic markers were similarly LE-enriched (Supp. Material), consistent with the contribution of resident brain cells to this compartment. At the same time, *L1CAM*, a mediator of GBM migration along white matter tracts [63], emerged as a strong LE signal, indicating that the leading-edge profile is not explained solely by contamination from normal tissue but also captures biology associated with tumor invasion. Together, these patterns distinguish tumor-intrinsic programs concentrated in the cellular tumor from transcriptional signals arising at the interface between infiltrating malignant cells and surrounding brain tissue.

We next compared `TIDEST` with four published methods across the complete 90-gene panel, fitting each method independently to all 26 GBM sections and combining section-specific estimates using DerSimonian-Laird random-effects meta-analysis (BH *q <* 0.05) (Fig. **5e**). `TIDEST` recovered approximately 1.5–2× more significant genes than competing approaches while maintaining comparable direction accuracy among significant calls. `DESpace` and `SpatialGEE`, which explicitly model spatial structure, modestly improved direction accuracy over the naive *t*-test and `SpaGCN`, further underscoring the importance of spatial structure in this setting.

A notable feature of the data was that two classes of genes previously predicted from single-cell studies to be LE-enriched were instead consistently CT-enriched in the Visium data (Fig. **5f**) [61]. The first comprised OPC-like and NPC-like GBM state markers, including *ASCL1* (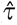 = 0.111, *q* = 8.9 10^−15^, significant in 88% of samples), *OLIG1* (*q* = 4.4 10^−11^, 81%), *OLIG2* (*q* = 8.2 10^−11^, 73%), *SOX10* (*q* = 5.0 10^−7^, 77%), and *PDGFRA* (*q* = 2.6 10^−8^, 58%). The second comprised invasion-associated extracellular matrix genes, including *BCAN* (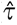 = 0.195, *q* = 3.9 10^−13^, 92%) and *PTPRZ1* (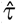 = 0.119, *q* = 1.6 10^−9^, 88%). All four competing methods recovered the same CT-enriched direction for all seven genes (Supp. Table D.3), arguing against a method-specific artifact. Instead, the discrepancy likely reflects the 55-*μ*m resolution of Visium spots, which can mix tumor cells across spatial compartments. This observation distinguishes inferential uncertainty from measurement resolution: accounting for reconstruction error and spatial confounding can improve inference, but cannot recover cellular resolution absent from the underlying measurement.

### TIDEST redefines the transcriptional landscape of human breast cancer invasion

Progression from ductal carcinoma *in situ* (DCIS) to invasive breast carcinoma (IBC) is accompanied by extensive remodeling of tissue architecture, including disruption of the myoepithelial barrier, activation of stromal fibroblasts, and immune infiltration. Consequently, spatial expression differences between pre-invasive and invasive regions reflect not only malignant progression but also substantial changes in local cellular composition, creating a setting in which latent spatial variation can alter the inferred transcriptional programs of invasion.

We applied `TIDEST` to a Xenium single-cell spatial transcriptomics dataset of human breast tissue containing both DCIS and IBC within the same tissue section [3], analyzing 62755 annotated locations (Fig. **6a**). We defined *A_i_* = 1 for IBC and *A_i_* = 0 for DCIS, so positive estimates correspond to higher expression in invasive regions. Because the Xenium panel measures only 313 genes, we reconstructed the transcriptome using `CellPLM` [21], a pre-trained cell language model, together with a matched scRNA-seq reference comprising 27000 cells [3], and used the reconstructed expression to construct the augmented outcomes.

**Figure 6:**
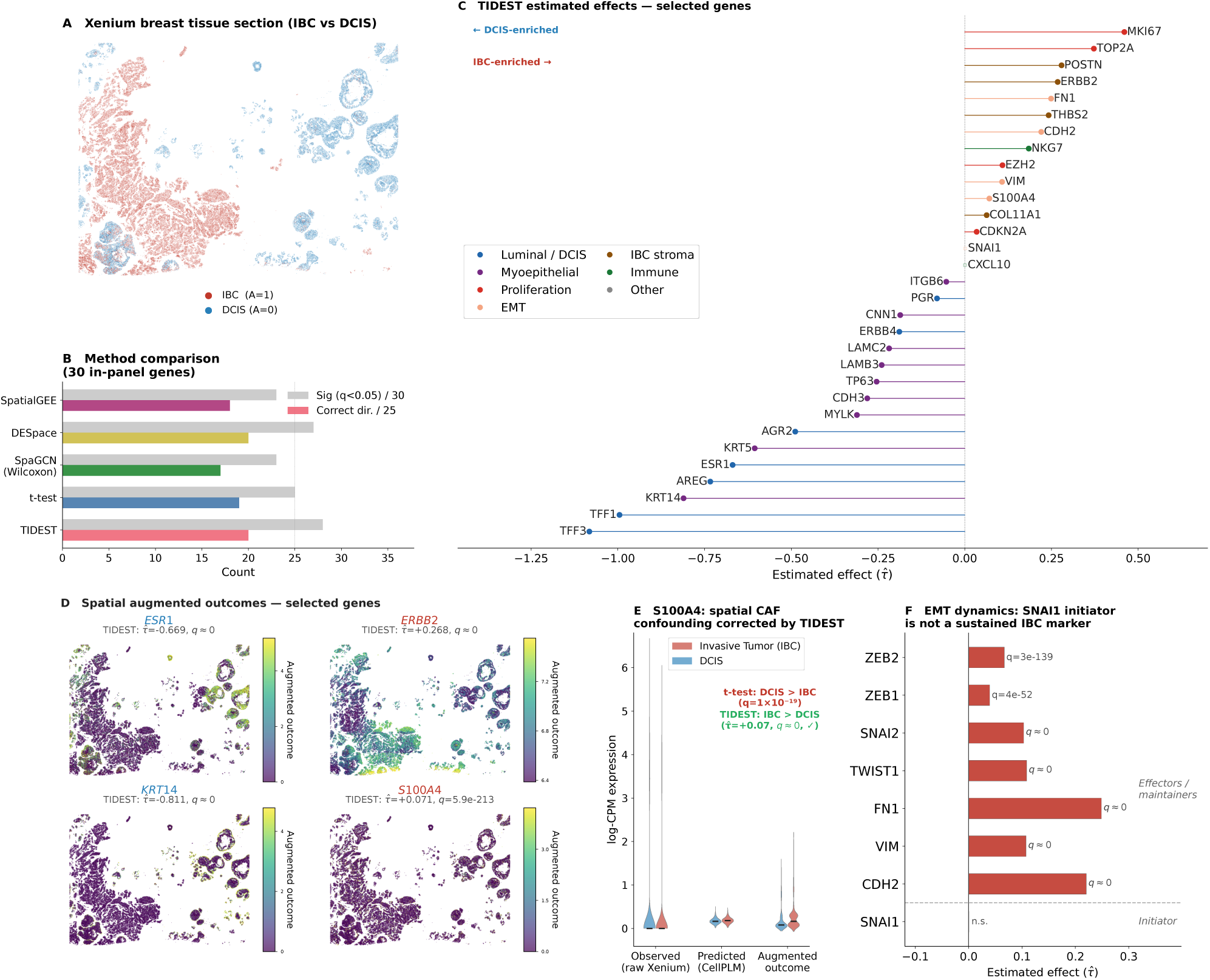
Human breast cancer application. **(a)** Xenium spatial transcriptomics section of human breast tissue (62755 cells). Each point is a single cell; red = IBC (*A* = 1, *n* = 28961); blue = DCIS (*A* = 0, *n* = 33794). **(b)** Method comparison on the 30 genes present in the observed Xenium panel. Grouped horizontal bars show, for each method, the number of genes called significant out of 30 (gray) and the number that are both significant and correctly directioned out of the 25 directional in-panel genes (red, `TIDEST` color). Dashed vertical line marks 25 (total directional genes). `TIDEST` and `DESpace` tie at 20/25 correct directions. **(c)** `TIDEST` estimated effects 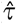 for selected labeled marker genes, sorted by effect size (DCIS-enriched at left, IBC-enriched at right). Color encodes biological group (see legend). Dashed vertical line at 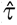 = 0. **(d)** Spatial augmented outcome maps for four key genes (*ESR1*, *ERBB2*, *KRT14*, *S100A4*). Gene name color encodes direction (red = IBC-enriched; blue = DCIS-enriched). Subtitle reports the `TIDEST` 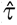 and *q*-value. **(e)** *S100A4*: distributions of observed Xenium expression (left), `CellPLM` predicted expression (center), and augmented outcome (right), split by IBC (red) and DCIS (blue). Naive comparison using *t*-test on observed expression returns DCIS *>* IBC (*q* = 1 × 10^−19^, incorrect direction). `TIDEST` on the augmented outcome recovers the biologically expected IBC enrichment (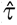 = +0.07, *q* ≈ 0). **(f)** EMT dynamics. *SNAI1* (initiator) is non-significant (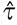 ≈ 0, *q* = 0.15); all seven downstream EMT effectors and structural remodeling markers are significantly IBC-enriched, consistent with a tumor that has completed rather than is currently undergoing mesenchymal transition.

Accounting for latent spatial variation altered the biological interpretation of several genes associated with invasion. The clearest example was the cancer-associated fibroblast marker *S100A4*, which has an established role in tumor invasion and is expected to be enriched in IBC [64]. The naive *t*-test identified *S100A4* as significantly differentially expressed in the opposite direction, with higher expression in DCIS. This reversal is consistent with fibroblasts interdigitated around DCIS ducts contributing disproportionately to local cellular composition and thereby biasing conventional comparisons. After adjustment with `TIDEST`, *S100A4* was recovered as IBC-enriched (Fig. **6e**), demonstrating that latent spatial variation can invert the inferred direction of differential expression for biologically important invasion-associated genes. *NKG7* provides a second example of this phenomenon (Supp. Material).

`TIDEST` also refined the interpretation of regulatory programs governing invasion. *SNAI1*, a canonical initiator of epithelial-to-mesenchymal transition (EMT) that represses *CDH1* to induce cellular plasticity [65], was not significantly differentially expressed between DCIS and IBC (Fig. **6f**). In contrast, downstream components of the EMT transcriptional network – including *CDH2*, *VIM*, *TWIST1*, *FN1*, *ZEB1*, and *ZEB2* – were all significantly enriched in invasive regions. This pattern distinguishes transient activation of an upstream regulatory factor from persistent activation of its downstream transcriptional program, consistent with an established invasive phenotype in which mesenchymal gene expression is maintained after *SNAI1* expression has subsided. Rather than indicating ongoing EMT induction, these data suggest that the invasive regions have progressed to a stable mesenchymal transcriptional state.

These changes in biological interpretation occurred within a broader transcriptional program consistent with progression from pre-invasive to invasive disease. We assembled a panel of 68 genes representing luminal differentiation, HER2 signaling, proliferation, EMT, extracellular matrix remodeling, immune infiltration, and myoepithelial integrity; 25 genes had well-established expected directionality in the IBC-versus-DCIS comparison. `TIDEST` identified 65 of 68 genes as significantly differentially expressed (BH *q <* 0.05) (Fig. **6b**). DCIS was enriched for canonical luminal differentiation genes, including *TFF3*, *TFF1*, *AREG*, *AGR2*, and the estrogen receptor *ESR1*. In contrast, IBC exhibited coordinated activation of proliferative, oncogenic, and invasive programs, including *MKI67*, *TOP2A*, the breast cancer driver *ERBB2*, the extracellular matrix regulator *POSTN*, and the mesenchymal marker *FN1* (Fig. **6c,d**) [66].

We next compared `TIDEST` with competing methods on the 30 genes directly measured by the Xenium panel (Fig. **6b**). Among the 25 directional markers, `TIDEST` recovered 20 with the expected direction, matching the best competing method while detecting the largest number of significant genes overall. Thus, the principal advantage of `TIDEST` in this setting was not simply increased detection, but the ability to distinguish transcriptional changes associated with invasion from differences arising from the spatial organization of the surrounding tissue.

Histological context also clarified the apparently paradoxical behavior of *KRT5* and *KRT14*, which were strongly enriched in DCIS despite frequently being described as basal-like breast cancer markers. In this spatial comparison, their expression is consistent with preservation of the myoepithelial layer surrounding DCIS ducts, which is lost following basement membrane invasion [67, 68]. This interpretation is supported by concordant enrichment of additional myoepithelial markers and was consistently recovered across competing methods (Supp. Material). Thus, apparent differences between pre-invasive and invasive regions can encode tissue architecture as well as tumor-cell state, reinforcing the importance of histological context when interpreting spatial differential expression.

## Discussion

Spatial transcriptomics is entering a new phase in which biological discovery increasingly depends on computationally reconstructed rather than directly measured transcriptomes. As reconstruction methods continue to improve, the challenge increasingly extends beyond prediction accuracy to the validity of downstream biological inference. Our results suggest that the next challenge is no longer reconstructing missing molecular measurements, but drawing valid biological conclusions from them. By treating reconstructed expression as experimentally observed, current analyses overlook both prediction uncertainty and the spatial organization of tissues, potentially producing misleading differential-expression signals and obscuring disease-associated transcriptional programs. This disconnect between prediction and inference represents an emerging challenge for computational spatial biology.

Across four biologically distinct systems, statistically valid inference consistently changed the interpretation of reconstructed spatial transcriptomes. In the mouse neocortex, it refined laminar transcriptional profiles by reducing biologically implausible differential-expression signals. In the human dorsolateral prefrontal cortex, it revealed a consistent convergence of genetic risk for educational attainment, intelligence and schizophrenia within Layer 2/3. In glioblastoma, it separated tumor-intrinsic transcriptional programs from the surrounding brain microenvironment at the invasive margin. In breast cancer, it redefined transcriptional programs associated with invasion by correcting misleading differential-expression calls and distinguishing persistent mesenchymal programs from transient regulatory activation. Together, these findings demonstrate that improving statistical inference does more than reduce false discoveries – it changes biological interpretation.

Several limitations warrant consideration. First, `TIDEST` depends on the availability of an appropriate single-cell reference, and performance may decline when the reference fails to capture the full cellular diversity represented in the spatial dataset. Second, the augmentation procedure assumes that prediction errors can be partially inferred from highly correlated observed genes; this assumption may be less reliable in settings with weak gene–gene correlation structure. Third, the current framework focuses on binary differential expression testing. Extending the methodology to continuous exposures, multiple experimental conditions, temporal trajectories, and hierarchical study designs represents an important direction for future research. Finally, variation in cell-type composition is a major determinant of spatial expression heterogeneity and may confound differential expression analyses. In the current implementation, such effects are absorbed into the estimated latent factors, enabling adjustment for broad compositional gradients without explicitly modeling cell identity. Nevertheless, integrating cell-type-specific information directly into the inferential framework may further improve power and interpretability.

Although developed for differential expression, the principles underlying `TIDEST` extend beyond this application. Prediction-powered calibration and statistically valid post-prediction inference may prove broadly useful for analyses of reconstructed molecular measurements, including gene-set enrichment, co-expression analysis and spatially variable gene detection. As computational reconstruction becomes routine across spatial omics, the central challenge will increasingly shift from predicting molecular measurements to interpreting them reliably. We anticipate that statistically valid inference will become an essential component of computational spatial biology, enabling increasingly accurate reconstruction to translate into reproducible biological discovery.

## Methods

### Details on TIDEST

*Testing for Imputed Differential Expression in Spatial Transcriptomics* (`TIDEST`) is a multi-stage framework designed to perform rigorous, spatially aware hypothesis testing on imputed spatial transcriptomics data. The pipeline addresses the dual challenges of prediction uncertainty – where downstream workflows naively treat noisy, deep learning-generated imputations as ground truth observations – and spatial autocorrelation, which violates standard sample independence assumptions and induces severe false-positive inflation in traditional statistical tests. To restore valid inference, `TIDEST` processes imputed expression matrices through three sequential modules: neighborhood augmentation debiasing, spatial gradient decomposition via surrogate-control outcomes, and orthogonalized DE effect estimation using partially linear models (PLMs).

To calibrate the systematic biases inherent in deep learning-based spatial reconstructions, `TIDEST` constructs a debiased augmented outcome *Ŷ_ij_* for each spatial coordinate *i* and target gene *j*. Rather than conducting downstream inference directly on the raw imputed values *Ỹ_ij_*, the framework leverages the empirical prediction residuals from a designated genomic neighborhood N_*j*_ consisting of highly correlated, physically measured anchor genes. For a given target gene *j*, N_*j*_ is populated by the most highly correlated, spatially observed genes (default: top 5 genes), where the feature-feature correlations 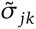 are estimated from a scRNA-seq reference matching the biological system. Let *Y_ik_* denote the true observed expression of neighbor gene *k ∊ N*_*j*_ at location *i*, and let *Ỹ_ik_* be its corresponding model imputation. The debiased augmented outcome for the target gene is defined as:

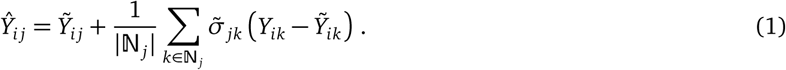

By averaging the measured errors of observed spatial anchors, this procedure dynamically reduces the bias of unmeasured or sparsely captured transcripts exploiting information from ground-truth signals.

Spatially resolved transcriptomics data exhibit pervasive spatial autocorrelation, where large-scale tissue architectures (e.g., anatomical layering or tumor-microenvironment gradients) create continuous spatial patterns. When these structural variations correlate with a binary condition or tissue boundary, they act as unobserved confounders *U*, inflating Type-I error rates by violating standard independence assumptions. `TIDEST` addresses this by implementing an empirical spatial whitening procedure anchored on surrogate-control outcomes [69]. This strategy is closely related to surrogate variable analysis, where latent factors estimated from control features are used to account for unobserved variation [29]. In the present setting, these latent factors may represent spatial structure, cellular composition, technical artifacts, or other unmeasured sources of heterogeneity. To safely isolate local biological variation from confounding tissue-level trends, `TIDEST` scans the transcriptomic panel to identify a set of surrogate-control genes that satisfy two precise operational constraints:

- **Negative control outcome.** The empirical Pearson correlation between the target genes of interest and the candidate surrogate controls computed on the scRNA-seq reference falls below a threshold *r*_NCO_.
- **Negative control exposure.** The spatial variability of the candidate surrogate controls – quantified by their coefficient of variation, defined as 100 * γ*/μ*, where γ is the standard deviation and *μ* is the mean of a given spatial gene expression – falls below a threshold *r*_NCE_.

These joint conditions guarantee that the selected surrogate-control genes are empirically decoupled from both the target biological outcomes and the primary treatment exposure. Once this control set is established, its spatial expression profile is utilized to reconstruct a low-dimensional representation of the latent confounding field *Û*. By default, `TIDEST` utilizes `SpatialPCA` to extract these latent principal components via a kernelized eigendecomposition [30], though alternative spatial dimension reduction techniques can be seamlessly integrated into the pipeline (e.g., `StaNMF` [70]).

In the final stage, `TIDEST` decouples the true treatment effect from large-scale spatial dependencies by mapping the augmented outcomes into a Robinson partially linear model (PLM) [31]. The conditional expectation of the debiased outcome is modeled as:

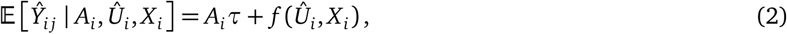

where *τ* represents the core estimand of interest (the treatment effect of the binary condition *A*), *Û* is the matrix of estimated confounders, and *X* captures additional spot-level covariates of interest (e.g., log-transformed library size). *f* (·) represents a flexible and unknown function mapping the spatial and technical confounders to expression levels.

By default, `TIDEST` utilizes random forests within a cross-fitting framework to flexibly estimate the nuisance functions in Equation (2). Crucially, this formulation yields an orthogonalized score function, guaranteeing that 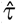 achieves asymptotic normality and valid confidence intervals even when the non-parametric nuisance estimators or latent spatial components are estimated at slower, non-parametric rates [69].

### Simulation study design

We generated synthetic datasets consisting of *n* = 300 spots on a regular two-dimensional grid, *p* = 200 genes partitioned into 10 non-overlapping co-expression modules (20 genes each). We generated the binary treatment *A_i_* using the “two-region” DGP: treatment is a clean rectangular split – left half of the grid is assigned with *A* = 0, right half has *A* = 1. This is the most adversarial setting for naive methods because the spatial confounder *U* (a smooth Gaussian field) correlates strongly with the treatment boundary. In the Supp. Material, we explore additional DGPs. The true latent log-expression for gene *j* at spot *i* follows

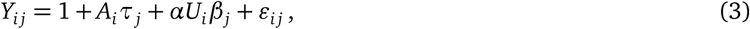

where 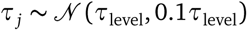 for the *p*_DE_ = 50 differentially expressed genes, with random sign, and *τ_j_* = 0 for the *p* - *p*_DE_ = 150 null genes; 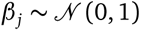 are gene-specific confounder loadings; 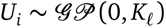 is a spatially smooth Gaussian random field with length-scale A = 0.3 that is independent of *A*; α ∊ *{*0, 0.25, 0.50, 0.75, 1.0*}* scales the confounder contribution; and 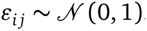. Observed counts are drawn as *C_ij_* ⇐ Poisson(exp(*Y_ij_*)).

The imputation model is parameterized as

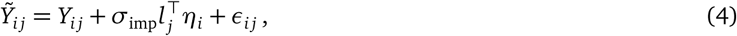

where 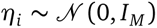 introduces module-structured noise, *l _j_* is the module loading vector of gene *j* (one entry is drawn from Uniform(0.5, 1.5), measuring the strength of its module membership; all the other entries are zero), and σ_imp_ 0.1, 0.5, 1.0, 1.5 parameterizes imputation quality (smaller means better). We also generate a synthetic single-cell reference of 2000 cells sharing the same module structure, so the Pearson correction exploits a realistic gene-gene correlation matrix.

We compared TIDEST against four spatial DE methods, all receiving identical observed count matrices and spatial coordinates: *t*-test (Welch two-sample *t*-test on log1p-normalized counts); SpaGCN [24] (Wilcoxon rank-sum test between spatial domains); DESpace [25] (negative-binomial model with spatial domain as a factor); and SpatialGEE [26] (generalized estimating equation with Poisson family and *k*-means working correlation structure).

In `TIDEST` pipeline, unobserved confounders were estimated using `SpatialPCA` [30] by truncated eigendecomposition of the Gaussian spatial kernel matrix retaining the top 20 eigenvectors. Eigenvectors with absolute Pearson correlation *r >* 0.5 with the treatment *A* were dropped from the confounder matrix *U* prior to fitting. The PLM was fitted with two cross-fitting folds and 100 regression trees per nuisance estimator.

The main simulation grid crossed five confounder levels α ∊ *{*0, 0.25, 0.50, 0.75, 1.0*}* with three DGP variants and fixed *τ*_level_ = 1.0, σ_imp_ = 0.5, yielding 15 settings; each was replicated 50 times. The DGP illustration panels (**A**-**D**) used *n* = 400 spots on a hexagonal grid for visual clarity. The augmentation sweep (panel **E**) varied σ_imp_ independently across six levels 0.1, 0.3, 0.5, 0.7, 1.0, 1.5 with 100 replicates per level, holding α = 0 to isolate reconstruction quality from confounding. Extended results including all α levels and DGP-variant breakdowns are provided in the Supp. Material.

### Real-data processing

#### Mouse brain data

We used a mouse neocortex Visium section dataset [33], restricting to spots annotated as “Cortex_1” (deep layers L5/6, *n* = 171) or “Cortex_3” (superficial layers L2/3/4, *n* = 126; 297 spots total). We assembled a panel of 55 curated layer-marker genes comprising 15 genes expected to be superficial-enriched, 23 deep-enriched, and 17 additional markers (including cell-type markers and negative controls); expected directions were assigned from published layer-specific expression atlases [35, 71]. The binary treatment is *A_i_* = 1 for superficial spots and *A_i_* = 0 for deep spots, so a positive *τ* indicates higher superficial expression. The scRNA-seq reference was an adult mouse cortex dataset [34] (21697 cells), normalized to 10^4^ counts. `Tangram` [10] was fitted with RNA-count-based density prior; imputed expression was Pearson-augmented using the top-5 nearest-neighbors. `SpatialPCA` [30] was run retaining 10 principal components; one PC with absolute correlation *|ρ|* = 0.779 with the treatment *A* was dropped by the correlation screen (threshold *ρ >* 0.5), leaving nine `SpatialPCA` components as an estimate of unmeasured confounding. Log library size was appended as a covariate. The Robinson PLM was fitted with two cross-fitting folds and 200 regression trees per nuisance estimator (random-forest implementation). The four competing methods (*t*-test, `SpaGCN` Wilcoxon, `DESpace`, and `SpatialGEE`) were applied with identical settings to the simulation study, operating on the log1p-normalized observed Visium counts. All methods received the same 55 marker genes, and FDR control was applied independently within each method using the Benjamini-Hochberg procedure.

#### Human brain data

We analyzed 12 dorsolateral prefrontal cortex (DLPFC) 10x Visium spatial transcriptomics sections from three neurotypical donors [48]. Each section is annotated with manual cortical-layer labels; we merged Layer 2 and Layer 3 into a single “Layer 2/3” category for consistency across sections, retained only spots annotated as Layer 1, Layer 2/3, Layer 4, Layer 5, or Layer 6, and discarded spots annotated as white matter. For each of three GWAS traits under study – educational attainment (EA) [49], schizophrenia (SCZ) [50], and intelligence (IQ) [51] – we queried the Open Targets Platform Locus-to-Gene (L2G) pipeline for the top-predicted gene at each GWAS credible set, yielding 200 (EA), 221 (SCZ), and 146 (IQ) trait genes. Each trait gene was paired with 10 expression-matched background candidate genes (nearest neighbors in pooled observed log-CPM expression across the 12 sections, restricted to genes retained by `Tangram` in every sample), giving background panels of 2000 (EA), 2210 (SCZ), and 1460 (IQ) genes tested alongside the corresponding trait genes. As `TIDEST` requires a single-cell reference for imputation, we used a DLPFC snRNA-seq reference [72] (5231 nuclei from 2 neurotypical donors, disjoint from the 10x Visium cohort). The single-cell reference was normalized to 10^4^ counts. For `Tangram` [10] training we selected the top 100 marker genes per reference cell type (including layer-resolved excitatory subtypes), using the reference cell-type labels as the grouping variable. `Tangram` was fitted in `cells` mode with an RNA-count-based density prior.

The augmented outcome for each spot was constructed using the Pearson correlation correction with top-5 neighbors, with the Pearson correlation matrix computed from the snRNA-seq reference. We applied per-spot total-count scaling to align the `Tangram` imputed expression to the observed 10x Visium count scale before log-transformation. For each sample, we selected confounder genes for `SpatialPCA` as those with coefficient of variation *r_NCE_ <* 0.3 in the augmented outcomes and Pearson correlation *|r_NCO_| <* 0.3 with any panel gene in the scRNA-seq reference. The top 2000 such genes ranked by spatial variance were used to fit `SpatialPCA` with 10 principal components. For each sample, `SpatialPCA` PCs with correlation with the treatment greater than 0.5 were excluded from the confounder matrix before fitting the PLM. Log-library size was appended as an additional covariate. For each of the 5 cortical layers, we fit the PLM independently for each of the 12 sections – treatment defined as a spot being annotated to that layer versus all other retained layers – using 2-fold cross-fitting and random forests with 200 trees. Per-sample *p*-values were adjusted by the Benjamini-Hochberg procedure within each sample (*q <* 0.05).

Cross-sample meta-analysis used DerSimonian-Laird random-effects inverse-variance weighting to account for between-sample heterogeneity, followed by a second round of Benjamini-Hochberg correction over all genes tested within that layer. The between-sample variance 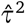 was estimated as 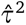 = max 0, (*Q* − (*k* − 1))*/c*, where 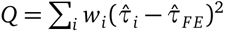 is Cochran’s heterogeneity statistic, *k* = 12, and 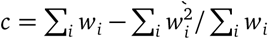 with 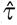. To assess the stability of the layer ranking by enrichment, we performed a sample-level bootstrap (2000 replicates). In each replicate, for every cortical layer we resampled the 12 sections with replacement and recomputed the DerSimonian-Laird random-effects meta-analysis (as above) on the resampled sample set. Restricting to genes with positive 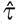_*RE*_, we computed a rank-based effect size – the common language effect size (CLES), the probability that a randomly drawn trait gene’s 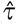_*RE*_ exceeds that of a randomly drawn background gene, obtained from the Mann-Whitney *U* statistic – comparing the trait genes to all background candidate genes within that layer. The 5 layers were then ranked by this CLES value (rank 1 = most enriched, rank 5 = least enriched) for that replicate. Repeating this across all 2000 replicates gives, for each layer, an empirical distribution over its rank position, which we use to quantify how consistently each layer is identified as the most (or least) trait-enriched under resampling of the 12 sections.

#### Human glioblastoma data

We analyzed 26 IDH-wildtype GBM Visium spatial transcriptomics sections from three patient cohorts [57]: the Massachusetts General Hospital (MGH) cohort (*n* = 1), the University of Freiburg (UKF) cohort (*n* = 13), and the University Hospital Zurich (ZH) cohort (*n* = 12). Each section is annotated with IvyGAP histological regions [58]; we retained only spots annotated as cellular tumor (CT) or leading edge (LE), discarding spots with other IvyGAP annotations. We assembled a curated panel of 90 genes (predicted) representative of the major GBM molecular programs, normal brain cell types, immune microenvironment, and metabolic states. As `TIDEST` requires a single-cell reference for imputation, we used the human GBM scRNA-seq dataset [59] (3589 cells from a disjoint cohort of patients). The single-cell reference was normalized to 10^4^ counts. For `Tangram` [10] training we selected the top 100 marker genes per reference cell-type, using the reference cell-type labels as the grouping variable. `Tangram` was fitted in `cells` mode with an RNA-count-based density prior.

The augmented outcome for each spot *i* was constructed using the Pearson correlation correction with top-5 neighbors. The Pearson correlation matrix was computed once from the scRNA-seq reference and shared across all 26 samples. We applied per-spot total-count scaling to align the `Tangram` imputed expression to the observed Visium count scale before log-transformation. For each sample, we selected confounder genes for `SpatialPCA` as those with coefficient of variation *r_NCE_ <* 0.5 in the augmented outcomes and Pearson correlation *|r_NCO_| <* 0.3 with any panel gene in the scRNA-seq reference. The top 2000 such genes ranked by spatial variance were used to fit `SpatialPCA` with 50 principal components. For each sample, `SpatialPCA` PCs with correlation with the treatment greater than 0.5 were excluded from the confounder matrix before fitting the PLM. Log-library size was appended as an additional covariate. We fitted the PLM independently for each sample using 5-fold cross-fitting and random forests with 200 trees. Per-sample *p*-values were adjusted by the Benjamini-Hochberg procedure within each sample (*q <* 0.05). Cross-sample meta-analysis used DerSimonian-Laird random-effects inverse-variance weighting (as above) to account for between-tumor heterogeneity, with *k* = 26, followed by a second BH correction over the 90 panel genes.

#### Human breast cancer data

We used a Xenium FFPE spatial transcriptomics section of human breast tissue [3] containing both invasive breast cancer and ductal carcinoma *in situ* in the same specimen (62755 cells, 313-gene panel). Cluster annotations were taken from the companion single-cell reference [3]. The binary treatment was defined as *A_i_*= 1 for cells annotated as “Invasive_Tumor” or “Prolif_Invasive_Tumor” (IBC, *n* = 28961) and *A_i_* = 0 for cells annotated as “DCIS_1” or “DCIS_2” (DCIS, *n* = 33794). We assembled a panel of 68 curated marker genes spanning IBC proliferation and invasion drivers, DCIS luminal markers, myoepithelial keratins, EMT effectors and maintainers, immune infiltrate markers, and contested genes; 25 genes have a well-established expected direction in the IBC-versus-DCIS contrast, assigned from published IBC, DCIS, and myoepithelial expression studies [67, 73, 74]. The expected direction for *KRT5* and *KRT14* was set to DCIS, as these myoepithelial markers reflect the intact basement membrane present in DCIS rather than the basal subtype designation used in gene-panel studies. Genes with unclear expected directions due to ER/HER2-status dependence (e.g., *EGFR*, *FOXA1*, *MLPH*, *GATA3*) were excluded from the directional evaluation.

Because the Xenium panel covers only 313 genes, we constructed the augmented outcome using `CellPLM` [21], a pre-trained Transformer-based cell language model. The matched single-cell reference (27000 cells, 18085 genes) was normalized to 10^4^ counts per cell and log1p-transformed before input to `CellPLM`; the same normalization was applied to the Xenium ST data. `CellPLM` outputs expression in log-CPM space, so the Pearson correction is applied directly as the difference between observed and predicted log-CPM, without additional scaling. The top-5 nearest-neighbor Pearson correction was applied to construct the augmented outcome. Confounder genes were selected using coefficient of variation threshold *r_NCE_ <* 1.5 and marker correlation threshold *|r_NCO_|* = 0.3, retaining 2000 genes; the higher *r_NCE_* threshold (compared to the Visium applications) reflects the greater single-cell-level expression variability in Xenium data.

Due to the large cell count, fitting `SpatialPCA` directly would require constructing a 62755 × 62755 kernel matrix, which exceeds available memory. We instead fitted `SpatialPCA` on a stratified subsample of 10000 cells (5000 per group), retaining 50 principal components, and then projected all 62755 cells into the fitted `SpatialPCA` space. No `SpatialPCA` component with absolute Pearson correlation *ρ >* 0.5 with the treatment *A* was found, so none was excluded from the confounder matrix; log library size (log-CPM total per cell) was appended as a covariate. The PLM was fitted with 2 cross-fitting folds and 200 regression trees per nuisance estimator, retaining 50 `SpatialPCA` components (after the correlation screen) as confounders. Sandwich standard errors were computed from the cross-fitted residuals.

The four competing methods were applied to the 30 genes present in the observed Xenium panel, operating on log-CPM-normalized Xenium counts. For `DESpace`, 13 cells with zero total counts across the marker panel were filtered before constructing the `SpatialExperiment` object, as the `calcNormFactors` step in `edgeR` produces undefined normalization factors for zero-library cells. For `SpatialGEE`, a stratified subsample of 20000 cells (10000 per group) was used. FDR control was applied within each method using the Benjamini-Hochberg procedure. `TIDEST` additionally tested the 38 genes accessible only through `CellPLM` augmentation; these were not evaluated for competitors.

## Author contributions

All authors conceived ideas and analysis approaches. L.T. retrieved and processed data, implemented pipelines and performed statistical analyses. All authors interpreted findings and participated to the writing of the manuscript. J.L. and K.R. supervised the research.

## Data and code availability

Mouse brain application data are available directly from the `squidpy` package [75] through the built-in functions `datasets.visium_fluo_adata_crop()` (10x Visium) and `datasets.sc_mouse_cortex()` (scRNA-seq). Human brain application spatial (10x Visium) data can be downloaded from the HumanPilot `GitHub` webpage at https://github.com/LieberInstitute/HumanPilot. Single-cell reference data for the human brain application are available at this `GitHub` link: https://github.com/LieberInstitute/10xPilot_snRNAseq-human. GWAS credible sets were retrieved using the Open Targets GraphQL API, accession numbers GCST006250 (intelligence), GCST90503210 (schizophrenia), and GCST90105038 (educational attainment). Human glioblastoma application spatial data (10x Visium) are available under the accession number GSE237183 (GEO) and also stored on Zenodo at https://doi.org/10.5281/zenodo.8105466. Single-cell reference data for the human glioblastoma application are available under the accession number GSE84465 (GEO). Human breast cancer application data are available under the accession code GSE243280 (GEO) and also stored on 10x Genomics website at https://www.10xgenomics.com/products/xenium-in-situ/preview-dataset-human-breast. All code to reproduce the entire analysis pipeline, including simulations and real-data applications, as well as code containing `TIDEST` package, is available at https://github.com/testalorenzo/tidest.

## Acknowledgments

The authors wish to thank Francesca Chiaromonte, Bernie Devlin, Jin-Hong Du, Edward Kennedy, Eric Tchetgen Tchetgen, Qi Xu, and Bin Yu for useful feedback. This project was supported by National Institute of Mental Health (NIMH) grant R01MH123184.

## References

[1] Sophia K Longo, Margaret G Guo, Andrew L Ji, and Paul A Khavari. Integrating single-cell and spatial transcriptomics to elucidate intercellular tissue dynamics. Nature Reviews Genetics, 22(10):627–644, 2021.

[2] Vivien Marx. Method of the year: spatially resolved transcriptomics. Nature methods, 18(1):9–14, 2021.

[3] Amanda Janesick, Robert Shelansky, Andrew D Gottscho, Florian Wagner, Stephen R Williams, Morgane Rouault, Ghezal Beliakoff, Carolyn A Morrison, Michelli F Oliveira, Jordan T Sicherman, et al. High resolution mapping of the tumor microenvironment using integrated single-cell, spatial and in situ analysis. Nature communications, 14(1):8353, 2023.

[4] Samuel G Rodriques, Robert R Stickels, Aleksandrina Goeva, Carly A Martin, Evan Murray, Charles R Vanderburg, Joshua Welch, Linlin M Chen, Fei Chen, and Evan Z Macosko. Slide-seq: A scalable technology for measuring genome-wide expression at high spatial resolution. Science, 363(6434):1463–1467, 2019.

[5] Kok Hao Chen, Alistair N Boettiger, Jeffrey R Moffitt, Siyuan Wang, and Xiaowei Zhuang. Spatially resolved, highly multiplexed rna profiling in single cells. Science, 348(6233):aaa6090, 2015.

[6] Eric Lubeck, Ahmet F Coskun, Timur Zhiyentayev, Mubhij Ahmad, and Long Cai. Single-cell in situ rna profiling by sequential hybridization. Nature methods, 11(4):360–361, 2014.

[7] Sheel Shah, Eric Lubeck, Wen Zhou, and Long Cai. In situ transcription profiling of single cells reveals spatial organization of cells in the mouse hippocampus. Neuron, 92(2):342–357, 2016.

[8] Chee-Huat Linus Eng, Michael Lawson, Qian Zhu, Ruben Dries, Noushin Koulena, Yodai Takei, Jina Yun, Christopher Cronin, Christoph Karp, Guo-Cheng Yuan, et al. Transcriptome-scale super-resolved imaging in tissues by rna seqfish+. Nature, 568(7751):235–239, 2019.

[9] Xiaoyu Li, Fangfang Zhu, and Wenwen Min. Spadit: diffusion transformer for spatial gene expression prediction using scrna-seq. Briefings in Bioinformatics, 25(6):bbae571, 2024.

[10] Tommaso Biancalani, Gabriele Scalia, Lorenzo Buffoni, Raghav Avasthi, Ziqing Lu, Aman Sanger, Neriman Tokcan, Charles R Vanderburg, Åsa Segerstolpe, Meng Zhang, et al. Deep learning and alignment of spatially resolved single-cell transcriptomes with tangram. Nature methods, 18(11):1352–1362, 2021.

[11] Tamim Abdelaal, Soufiane Mourragui, Ahmed Mahfouz, and Marcel JT Reinders. Spage: spatial gene enhancement using scrna-seq. Nucleic acids research, 48(18):e107–e107, 2020.

[12] Chen Shengquan, Zhang Boheng, Chen Xiaoyang, Zhang Xuegong, and Jiang Rui. stplus: a reference-based method for the accurate enhancement of spatial transcriptomics. Bioinformatics, 37(Supplement_1):i299–i307, 2021.

[13] Zixuan Cang and Qing Nie. Inferring spatial and signaling relationships between cells from single cell transcriptomic data. Nature communications, 11(1):2084, 2020.

[14] Noa Moriel, Enes Senel, Nir Friedman, Nikolaus Rajewsky, Nikos Karaiskos, and Mor Nitzan. Novosparc: flexible spatial reconstruction of single-cell gene expression with optimal transport. Nature protocols, 16(9):4177–4200, 2021.

[15] Xiaomeng Wan, Jiashun Xiao, Sindy Sing Ting Tam, Mingxuan Cai, Ryohichi Sugimura, Yang Wang, Xiang Wan, Zhixiang Lin, Angela Ruohao Wu, and Can Yang. Integrating spatial and single-cell transcriptomics data using deep generative models with spatialscope. Nature Communications, 14(1):7848, 2023.

[16] Kongming Li, Jiahao Li, Yuhao Tao, and Fei Wang. stdiff: a diffusion model for imputing spatial transcriptomics through single-cell transcriptomics. Briefings in Bioinformatics, 25(3), 2024.

[17] Eric D Sun, Rong Ma, and James Zou. Sprite: improving spatial gene expression imputation with gene and cell networks. Bioinformatics, 40(Supplement_1):i521–i528, 2024.

[18] Tianyu Liu, Yingxin Lin, Xiao Luo, Yizhou Sun, and Hongyu Zhao. Vista uncovers missing gene expression and spatial-induced information for spatial transcriptomic data analysis. bioRxiv, pages 2024–08, 2025.

[19] Tianyu Liu, Tinglin Huang, Wengong Jin, Tinyi Chu, Rex Ying, and Hongyu Zhao. sprefine denoises and imputes spatial transcriptomics with a reference-free framework powered by genomic language model. bioRxiv, pages 2025–04, 2025.

[20] Hongjia Liu, Huamei Li, Amit Sharma, Guoyan Tang, Zongyu Xie, Yunyao Shen, Qiong Li, Chen Gong, Xiao Sun, Kun Luo, et al. Cellmap: precision mapping of cellular landscape in spatial transcriptomics. Nucleic Acids Research, 54(1):gkaf1484, 2026.

[21] Hongzhi Wen, Wenzhuo Tang, Xinnan Dai, Jiayuan Ding, Wei Jin, Yuying Xie, and Jiliang Tang. CellPLM: Pre-training of cell language model beyond single cells. In The Twelfth International Conference on Learning Representations, 2024. URL https://openreview.net/forum?id=BKXvPDekud.

[22] Romain Lopez, Achille Nazaret, Maxime Langevin, Jules Samaran, Jeffrey Regier, Michael I Jordan, and Nir Yosef. A joint model of unpaired data from scrna-seq and spatial transcriptomics for imputing missing gene expression measurements. arXiv preprint arXiv:1905.02269, 2019.

[23] Guanao Yan, Shuo Harper Hua, and Jingyi Jessica Li. Categorization of 34 computational methods to detect spatially variable genes from spatially resolved transcriptomics data. Nature Communications, 16(1):1141, 2025.

[24] Jian Hu, Xiangjie Li, Kyle Coleman, Amelia Schroeder, Nan Ma, David J Irwin, Edward B Lee, Russell T Shinohara, and Mingyao Li. Spagcn: Integrating gene expression, spatial location and histology to identify spatial domains and spatially variable genes by graph convolutional network. Nature methods, 18(11):1342–1351, 2021.

[25] Peiying Cai, Mark D Robinson, and Simone Tiberi. Despace: spatially variable gene detection via differential expression testing of spatial clusters. Bioinformatics, 40(2):btae027, 2024.

[26] Yishan Wang, Chenxuan Zang, Ziyi Li, Charles C Guo, Dejian Lai, and Peng Wei. A comparative study of statistical methods for identifying differentially expressed genes in spatial transcriptomics. bioRxiv, 2025.

[27] Anastasios N Angelopoulos, Stephen Bates, Clara Fannjiang, Michael I Jordan, and Tijana Zrnic. Prediction-powered inference. Science, 382(6671):669–674, 2023.

[28] Ergan Shang, Yuting Wei, and Kathryn Roeder. Predicting the unseen: a diffusion-based debiasing framework for transcriptional response prediction at single-cell resolution. Proceedings of the National Academy of Sciences, 122(52):e2525268122, 2025.

[29] Jeffrey T Leek and John D Storey. Capturing heterogeneity in gene expression studies by surrogate variable analysis. PLoS genetics, 3(9):e161, 2007.

[30] Lulu Shang and Xiang Zhou. Spatially aware dimension reduction for spatial transcriptomics. Nature communications, 13(1):7203, 2022.

[31] Peter M Robinson. Root-n-consistent semiparametric regression. Econometrica: journal of the Econometric Society, pages 931–954, 1988.

[32] Yuhan Hao, Tim Stuart, Madeline H Kowalski, Saket Choudhary, Paul Hoffman, Austin Hartman, Avi Srivastava, Gesmira Molla, Shaista Madad, Carlos Fernandez-Granda, and Rahul Satija. Dictionary learning for integrative, multimodal and scalable single-cell analysis. Nature Biotechnology, 2023. doi: 10.1038/s41587-023-01767-y. URL 10.1038/s41587-023-01767-y.

[33] 10x Genomics, 2020. URL https://cf.10xgenomics.com/samples/spatial-exp/1.1.0/V1_Adult_Mouse_Brain_Coronal_Section_2/V1_Adult_Mouse_Brain_Coronal_Section_2_web_summary.html.

[34] Bosiljka Tasic, Zizhen Yao, Lucas T Graybuck, Kimberly A Smith, Thuc Nghi Nguyen, Darren Bertagnolli, Jeff Goldy, Emma Garren, Michael N Economo, Sarada Viswanathan, et al. Shared and distinct transcriptomic cell types across neocortical areas. Nature, 563(7729):72–78, 2018.

[35] Bradley J Molyneaux, Paola Arlotta, Joao RL Menezes, and Jeffrey D Macklis. Neuronal subtype specification in the cerebral cortex. Nature reviews neuroscience, 8(6):427–437, 2007.

[36] Amit Zeisel, Hannah Hochgerner, Peter Lönnerberg, Anna Johnsson, Fatima Memic, Job Van Der Zwan, Martin Häring, Emelie Braun, Lars E Borm, Gioele La Manno, et al. Molecular architecture of the mouse nervous system. Cell, 174(4):999–1014, 2018.

[37] Tina Lai, Denis Jabaudon, Bradley J Molyneaux, Eiman Azim, Paola Arlotta, Joao RL Menezes, and Jeffrey D Macklis. Sox5 controls the sequential generation of distinct corticofugal neuron subtypes. Neuron, 57(2): 232–247, 2008.

[38] Kenneth Y Kwan, Mandy MS Lam, Željka Krsnik, Yuka Imamura Kawasawa, Veronique Lefebvre, and Nenad Šestan. Sox5 postmitotically regulates migration, postmigratory differentiation, and projections of subplate and deep-layer neocortical neurons. Proceedings of the National Academy of Sciences, 105(41):16021–16026, 2008.

[39] Xavier Caubit, Paolo Gubellini, Joris Andrieux, Pierre L Roubertoux, Mehdi Metwaly, Bernard Jacq, Ahmed Fatmi, Laurence Had-Aissouni, Kenneth Y Kwan, Pascal Salin, et al. Tshz3 deletion causes an autism syndrome and defects in cortical projection neurons. Nature genetics, 48(11):1359–1369, 2016.

[40] Bernardo Rudy, Gordon Fishell, SooHyun Lee, and Jens Hjerling-Leffler. Three groups of interneurons account for nearly 100% of neocortical gabaergic neurons. Developmental neurobiology, 71(1):45–61, 2011.

[41] Marta Nieto, Edwin S Monuki, Hua Tang, Jaime Imitola, Nicole Haubst, Samia J Khoury, Jim Cunningham, Magdalena Gotz, and Christopher A Walsh. Expression of cux-1 and cux-2 in the subventricular zone and upper layers ii–iv of the cerebral cortex. Journal of Comparative Neurology, 479(2):168–180, 2004.

[42] Bradley J Molyneaux, Paola Arlotta, Tustomu Hirata, Masahiko Hibi, and Jeffrey D Macklis. Fezl is required for the birth and specification of corticospinal motor neurons. Neuron, 47(6):817–831, 2005.

[43] Bin Chen, Song S Wang, Alexis M Hattox, Helen Rayburn, Sacha B Nelson, and Susan K McConnell. The fezf2–ctip2 genetic pathway regulates the fate choice of subcortical projection neurons in the developing cerebral cortex. Proceedings of the National Academy of Sciences, 105(32):11382–11387, 2008.

[44] Elizabeth A Alcamo, Laura Chirivella, Marcel Dautzenberg, Gergana Dobreva, Isabel Fariñas, Rudolf Grosschedl, and Susan K McConnell. Satb2 regulates callosal projection neuron identity in the developing cerebral cortex. Neuron, 57(3):364–377, 2008.

[45] Olga Britanova, Camino de Juan Romero, Amanda Cheung, Kenneth Y Kwan, Manuela Schwark, Andrea Gyorgy, Tanja Vogel, Sergey Akopov, Mišo Mitkovski, Denes Agoston, et al. Satb2 is a postmitotic determinant for upper-layer neuron specification in the neocortex. Neuron, 57(3):378–392, 2008.

[46] Z Josh Huang and Anirban Paul. The diversity of gabaergic neurons and neural communication elements. Nature Reviews Neuroscience, 20(9):563–572, 2019.

[47] Dino P Leone, Whitney E Heavner, Emily A Ferenczi, Gergana Dobreva, John R Huguenard, Rudolf Grosschedl, and Susan K McConnell. Satb2 regulates the differentiation of both callosal and subcerebral projection neurons in the developing cerebral cortex. Cerebral cortex, 25(10):3406–3419, 2015.

[48] Kristen R Maynard, Leonardo Collado-Torres, Lukas M Weber, Cedric Uytingco, Brianna K Barry, Stephen R Williams, Joseph L Catallini, Matthew N Tran, Zachary Besich, Madhavi Tippani, et al. Transcriptome-scale spatial gene expression in the human dorsolateral prefrontal cortex. Nature neuroscience, 24(3):425–436, 2021.

[49] Aysu Okbay, Yeda Wu, Nancy Wang, Hariharan Jayashankar, Michael Bennett, Seyed Moeen Nehzati, Julia Sidorenko, Hyeokmoon Kweon, Grant Goldman, Tamara Gjorgjieva, et al. Polygenic prediction of educational attainment within and between families from genome-wide association analyses in 3 million individuals. Nature genetics, 54(4):437–449, 2022.

[50] Xinglun Dang, Zhaowei Teng, Yongfeng Yang, Wenqiang Li, Jiewei Liu, Li Hui, Dongsheng Zhou, Daohua Gong, Shan-Shan Dai, Yifan Li, et al. Gene-level analysis reveals the genetic aetiology and therapeutic targets of schizophrenia. Nature Human Behaviour, 9(3):609–624, 2025.

[51] Jeanne E Savage, Philip R Jansen, Sven Stringer, Kyoko Watanabe, Julien Bryois, Christiaan A De Leeuw, Mats Nagel, Swapnil Awasthi, Peter B Barr, Jonathan RI Coleman, et al. Genome-wide association meta-analysis in 269,867 individuals identifies new genetic and functional links to intelligence. Nature genetics, 50(7):912–919, 2018.

[52] Lena Weißflog, Claus-Jürgen Scholz, Christian P Jacob, Thuy Trang Nguyen, Karin Zamzow, Silke Gro*β*-Lesch, Tobias J Renner, Marcel Romanos, Dan Rujescu, Susanne Walitza, et al. Kcnip4 as a candidate gene for personality disorders and adult adhd. European neuropsychopharmacology, 23(6):436–447, 2013.

[53] Winston H Dredge, Xuanjia Fan, Laura Sloofman, Lindsay Liang, Enrico Mossotto, Kendall Moore, Sarah Zipkowitz, Minghui Wang, Bin Zhang, Jiebiao Wang, et al. Spatiotemporal and genetic regulation of a-to-i editing throughout human brain development. Cell reports, 41(5), 2022.

[54] Mikihito Shibata, Kartik Pattabiraman, Sydney K Muchnik, Navjot Kaur, Yury M Morozov, Xiaoyang Cheng, Stephen G Waxman, and Nenad Sestan. Hominini-specific regulation of cbln2 increases prefrontal spinogenesis. Nature, 598(7881):489–494, 2021.

[55] Panfeng Zhao, Jian Yang, Jie Yang, and Jinfeng Du. Structural architecture and evolutionary conservation of cerebellin-mediated trans-synaptic signaling. Synapse, 80(3):e70045, 2026.

[56] Lloyd N Hopkins, Nesli Avgan, Heidi G Sutherland, Francesca E Fernandez, Emma EM Knowles, Larisa M Haupt, John Blangero, David C Glahn, David HK Shum, Rod A Lea, et al. Genome-wide association study identifies novel variants in olfactory, vitamin a, vitamin b, and cadherin pathways associated with learning and memory. Scientific Reports, 16(1):2911, 2025.

[57] Alissa C Greenwald, Noam Galili Darnell, Rouven Hoefflin, Dor Simkin, Christopher W Mount, L Nicolas Gonzalez Castro, Yotam Harnik, Sydney Dumont, Dana Hirsch, Masashi Nomura, et al. Integrative spatial analysis reveals a multi-layered organization of glioblastoma. Cell, 187(10):2485–2501, 2024.

[58] Ralph B Puchalski, Nameeta Shah, Jeremy Miller, Rachel Dalley, Steve R Nomura, Jae-Guen Yoon, Kimberly A Smith, Michael Lankerovich, Darren Bertagnolli, Kris Bickley, et al. An anatomic transcriptional atlas of human glioblastoma. Science, 360(6389):660–663, 2018.

[59] Spyros Darmanis, Steven A Sloan, Derek Croote, Marco Mignardi, Sophia Chernikova, Peyman Samghababi, Ye Zhang, Norma Neff, Mark Kowarsky, Christine Caneda, et al. Single-cell rna-seq analysis of infiltrating neoplastic cells at the migrating front of human glioblastoma. Cell reports, 21(5):1399–1410, 2017.

[60] Heidi S Phillips, Samir Kharbanda, Ruihuan Chen, William F Forrest, Robert H Soriano, Thomas D Wu, Anjan Misra, Janice M Nigro, Howard Colman, Liliana Soroceanu, et al. Molecular subclasses of high-grade glioma predict prognosis, delineate a pattern of disease progression, and resemble stages in neurogenesis. Cancer cell, 9 (3):157–173, 2006.

[61] Cyril Neftel, Julie Laffy, Mariella G Filbin, Toshiro Hara, Marni E Shore, Gilbert J Rahme, Alyssa R Richman, Dana Silverbush, McKenzie L Shaw, Christine M Hebert, et al. An integrative model of cellular states, plasticity, and genetics for glioblastoma. Cell, 178(4):835–849, 2019.

[62] Roel GW Verhaak, Katherine A Hoadley, Elizabeth Purdom, Victoria Wang, Yuan Qi, Matthew D Wilkerson, C Ryan Miller, Li Ding, Todd Golub, Jill P Mesirov, et al. Integrated genomic analysis identifies clinically relevant subtypes of glioblastoma characterized by abnormalities in pdgfra, idh1, egfr, and nf1. Cancer cell, 17(1):98–110, 2010.

[63] Matthias Osswald, Erik Jung, Felix Sahm, Gergely Solecki, Varun Venkataramani, Jonas Blaes, Sophie Weil, Heinz Horstmann, Benedikt Wiestler, Mustafa Syed, et al. Brain tumour cells interconnect to a functional and resistant network. Nature, 528(7580):93–98, 2015.

[64] Gil Friedman, Oshrat Levi-Galibov, Eyal David, Chamutal Bornstein, Amir Giladi, Maya Dadiani, Avi Mayo, Coral Halperin, Meirav Pevsner-Fischer, Hagar Lavon, et al. Cancer-associated fibroblast compositions change with breast cancer progression linking the ratio of s100a4+ and pdpn+ cafs to clinical outcome. Nature Cancer, 1(7): 692–708, 2020.

[65] Eduard Batlle, Elena Sancho, Clara Francí, David Domínguez, Mercè Monfar, Josep Baulida, and Antonio García de Herreros. The transcription factor snail is a repressor of e-cadherin gene expression in epithelial tumour cells. Nature cell biology, 2(2):84–89, 2000.

[66] Mahboubeh Sadeghi, Abbas Ghaderi, Pegah Mousavi, Soudabeh Sabetian, Amin Ramezani, and Mohammad Reza Haghshenas. Fn1 as a key gene in modulating the integrin cell surface pathway in breast cancer. *Medical oncology (Northwood, London*, England), 43(4):159, 2026.

[67] Werner Böcker, Roland Moll, Christopher Poremba, Roland Holland, Paul J Van Diest, Peter Dervan, Horst Bürger, Daniel Wai, Raihanatou Ina Diallo, Burkhard Brandt, et al. Common adult stem cells in the human breast give rise to glandular and myoepithelial cell lineages: a new cell biological concept. Laboratory investigation, 82(6): 737–746, 2002.

[68] Min Hu, Jun Yao, Li Cai, Kurt E Bachman, Frédéric Van Den Brûle, Victor Velculescu, and Kornelia Polyak. Distinct epigenetic changes in the stromal cells of breast cancers. Nature genetics, 37(8):899–905, 2005.

[69] Jin-Hong Du, Kathryn Roeder, and Larry Wasserman. Assumption-lean post-integrated inference with surrogate-control outcomes. Biometrika, page asag004, 2026.

[70] Siqi Wu, Antony Joseph, Ann S Hammonds, Susan E Celniker, Bin Yu, and Erwin Frise. Stability-driven nonnegative matrix factorization to interpret spatial gene expression and build local gene networks. Proceedings of the National Academy of Sciences, 113(16):4290–4295, 2016.

[71] Bosiljka Tasic, Vilas Menon, Thuc Nghi Nguyen, Tae Kyung Kim, Tim Jarsky, Zizhen Yao, Boaz Levi, Lucas T Gray, Staci A Sorensen, Tim Dolbeare, et al. Adult mouse cortical cell taxonomy revealed by single cell transcriptomics. Nature neuroscience, 19(2):335–346, 2016.

[72] Matthew N Tran, Kristen R Maynard, Abby Spangler, Louise A Huuki, Kelsey D Montgomery, Vijay Sadashivaiah, Madhavi Tippani, Brianna K Barry, Dana B Hancock, Stephanie C Hicks, et al. Single-nucleus transcriptome analysis reveals cell-type-specific molecular signatures across reward circuitry in the human brain. Neuron, 109 (19):3088–3103, 2021.

[73] Charles M Perou, Therese Sørlie, Michael B Eisen, Matt Van De Rijn, Stefanie S Jeffrey, Christian A Rees, Jonathan R Pollack, Douglas T Ross, Hilde Johnsen, Lars A Akslen, et al. Molecular portraits of human breast tumours. nature, 406(6797):747–752, 2000.

[74] Lily Xu, Kaitlyn Saunders, Shao-Po Huang, Hildur Knutsdottir, Kenneth Martinez-Algarin, Isabella Terrazas, Kenian Chen, Heather M McArthur, Julia Maués, Christine Hodgdon, et al. A comprehensive single-cell breast tumor atlas defines epithelial and immune heterogeneity and interactions predicting anti-pd-1 therapy response. Cell Reports Medicine, 5(5), 2024.

[75] Giovanni Palla, Hannah Spitzer, Michal Klein, David Fischer, Anna Christina Schaar, Louis Benedikt Kuemmerle, Sergei Rybakov, Ignacio L Ibarra, Olle Holmberg, Isaac Virshup, et al. Squidpy: a scalable framework for spatial omics analysis. Nature methods, 19(2):171–178, 2022.

