## Supplementary material for "Accurate prediction in reconstructed spatial transcriptomes does not ensure valid biological discovery": Supp. Material

#### A Theoretical framework for TIDEST

Here, we show that the confounder-adjustment and PLM-estimation stages of TIDEST can be seen as an extension of the assumption-lean post-integrated inference (PII) framework [1]. We then state and prove one result specific to TIDEST, that characterizes the conditions under which the neighborhood-augmentation step (Equation (1)) preserves the validity of this approach to inference. We then discuss the practical scope of these results.

##### A.1 Setup and target estimand

Fix a target gene $j$. For spot $i = 1, \ldots, n$, let $A_i \in \{0, 1\}$ denote the binary condition of interest, $X_i \in \mathbb{R}^q$ additional spot-level covariates (e.g., log library size), and $U_i \in \mathbb{R}^r$ an unobserved latent field summarizing spatial location, cellular composition, and other technical or biological sources of variation. Let $Y_{ij}$ denote the (unobserved) true expression of gene $j$ at spot $i$, related to $A_i, X_i, U_i$ through the partially linear model

$$\mathbb{E}\left[Y_{ij} \mid A_i, X_i, U_i\right] = A_i \bar{\tau}_j + f_j(X_i, U_i), \tag{5}$$

with $f_j$ an unrestricted function. The scientific estimand $\bar{\tau}_j$ – the quantity TIDEST aims to estimate and test – is identified as:

$$\bar{\tau}_j = (\mathbb{E}[\mathbb{V}[A \mid X, U)]])^{-1}\, \mathbb{E}\left[\mathrm{Cov}(A, Y_j \mid X, U)\right]. \tag{6}$$

TIDEST cannot evaluate Equation (6) directly because $Y_{ij}$ is not observed and thus replaced by the augmented outcome $\hat{Y}_{ij}$ (Equation (1)), and $U_i$ is unobserved and replaced by an embedding $\hat{U}_i$ estimated from data. Define the corresponding *feasible* estimand

$$\tau_j = \left(\mathbb{E}\left[\mathbb{V}\left[A \mid X, \hat{U})\right]\right]\right)^{-1} \mathbb{E}\left[\mathrm{Cov}(A, \hat{Y}_j \mid X, \hat{U})\right], \tag{7}$$

which is exactly the estimand targeted by Equation (2) in the Methods. Below we first relate $\hat{\tau}_j$ (the PLM output) to $\tau_j$, and then we relate $\tau_j$ to the scientific quantity $\bar{\tau}_j$.

##### A.2 Estimation and inference procedure for $\hat{\tau}_j$

The embedding $\hat{U}_i$ is constructed by TIDEST from the surrogate-control gene panel $\mathscr{C}$ (Methods, $r_{\mathrm{NCO}}/r_{\mathrm{NCE}}$ criteria): genes whose augmented expression is (a) approximately uncorrelated with the target genes in the SC reference ($r_{\mathrm{NCO}}$, negative control outcome) and (b) has low coefficient of variation ($r_{\mathrm{NCE}}$, negative control exposure). In the notation of [1], $\hat{Y}_{i,\mathscr{C}}$ plays the role of the *surrogate-control outcomes*: a subset of outcomes for which the direct effect of $A$ is, by construction of $\mathscr{C}$, approximately zero, so that their spatial variation is informative about $U$ and not about $\tau$. TIDEST then sets $\hat{U}_i = g_K(\hat{Y}_{i,\mathscr{C}})$, where $g_K$ is the SpatialPCA kernel eigenmap (top $K$ components after the $|\rho(\mathrm{PC}, A)| > 0.5$ correlation screen), playing the role of the embedding map of [1]. The correlation screen is an additional, TIDEST-specific safeguard against retaining components that are themselves strongly associated with $A$ (which would risk over-controlling, i.e., partialling out part of the treatment effect itself); it is not required by the theory below but is a conservative refinement of it (in case, by mistake, some active genes were inserted in the surrogate-control set).

Given $\hat{U}_i$ and $X_i$, TIDEST estimates $\tau_j$ in Equation (7) using the cross-fitted, double-residual procedure of [1], specialized to a scalar treatment $A$. With cross-fitting, for each fold we estimate out-of-fold regressions (default: random forests) $\hat{m} \approx \mathbb{E}\left[A \mid \hat{U}\right]$ and $\hat{\ell}_j \approx \mathbb{E}\left[\hat{Y}_j \mid \hat{U}\right]$, and we evaluate them in:

$$\hat{\tau}_j = \frac{\sum_{i=1}^{n}(A_i - \hat{m}(\hat{U}_i))(Y_{ij} - \hat{\ell}_j(\hat{U}_i))}{\sum_{i=1}^{n}(A_i - \hat{m}(\hat{U}_i))^2}. \tag{8}$$

Corollary 1, part 1, in [1] shows that (under assumptions) $\hat{\tau}_j$ converges in distribution to a Normal distribution centered in $\tau_j$. This establishes that, conditional on the embedding $\hat{U}$, TIDEST's PLM delivers asymptotically valid inference for the feasible estimand $\tau_j$ even though the nuisance functions $\hat{m}, \hat{\ell}_j$ are estimated nonparametrically (random forests) at slower-than-$\sqrt{n}$ rates – this is the orthogonality property invoked, but not previously stated, in the Methods. Two gaps remain between $\tau_j$ and the scientific estimand $\bar{\tau}_j$: the effect of replacing $U$ by $\hat{U}$, and the effect of replacing $Y_j$ by $\hat{Y}_j$.

The former gap is addressed again by [1]. In Corollary 1, part 2, it is shown that if $\|\hat{U} - U\|_2 = o_P(n^{-1/2})$, that is, if the learned embedding $\hat{U}$ converges to the true confounding field at parametric rate, then asymptotic normality still holds.

The remaining gap between the feasible estimand $\tau_j$ and the scientific estimand $\bar{\tau}_j$ is the substitution of the augmented outcome $\hat{Y}_j$ for the true expression $Y_j$. Write the *augmentation gap* as

$$\Delta_{ij} = \hat{Y}_{ij} - Y_{ij} = \underbrace{(\tilde{Y}_{ij} - Y_{ij})}_{\text{imputation error, gene } j} + \frac{1}{|\mathbb{N}_j|} \sum_{k \in \mathbb{N}_j} \tilde{\sigma}_{jk} \underbrace{(Y_{ik} - \tilde{Y}_{ik})}_{\text{imputation error, gene } k}, \tag{9}$$

directly from Equation (1). By linearity of covariance, $\text{Cov}(A, \hat{Y}_j \mid X, U) = \text{Cov}(A, Y_j \mid X, U) + \text{Cov}(A, \Delta_j \mid X, U)$, so

$$(\mathbb{E}[\mathbb{V}[A \mid X, U]])^{-1} \, \mathbb{E}\big[\text{Cov}(A, \hat{Y}_j \mid X, U)\big] = \bar{\tau}_j + (\mathbb{E}[\mathbb{V}[A \mid X, U]])^{-1} \, \mathbb{E}\big[\text{Cov}(A, \Delta_j \mid X, U)\big]. \tag{10}$$

**Lemma A.1** (Augmentation preserves the target estimand). *Suppose that (i) for every $k \in \{j\} \cup \mathbb{N}_j$, $\text{Cov}(\tilde{Y}_{ik} - Y_{ik}, A_i \mid X_i, U_i) = o_P(n^{-1/2})$, that is, the imputation residual for gene $k$ is asymptotically mean-independent of treatment given the confounders, and (ii) the neighbor weights $\tilde{\sigma}_{jk}$ are estimated from a reference dataset independent of $(A_i, X_i, U_i, Y_{i,k}, \tilde{Y}_{i,k})_{i=1}^n$. Then $\text{Cov}(A, \Delta_j \mid X, U) = o_p(n^{-1/2})$.*

*Proof.* By (9) and the triangle inequality,

$$\left|\text{Cov}(A, \Delta_j \mid X, U)\right| \le \left|\text{Cov}(A, \tilde{Y}_j - Y_j \mid X, U)\right| + \frac{1}{|\mathbb{N}_j|} \sum_{k \in \mathbb{N}_j} |\tilde{\sigma}_{jk}| \cdot \left|\text{Cov}(A, Y_k - \tilde{Y}_k \mid X, U)\right|,$$

using (ii) to treat $\tilde{\sigma}_{jk}$ as a fixed (data-independent) bounded weight ($|\tilde{\sigma}_{jk}| \le 1$ as a Pearson correlation). Each term is $o_p(n^{-1/2})$ by (i), and $|\mathbb{N}_j|$ is fixed, so the sum is $o_p(n^{-1/2})$. □

**Remark A.2.** Condition (i) does not require the imputation model to be accurate – no rate condition is placed on $\tilde{Y}_{ik} - Y^{\star}_{ik}$ itself. It only requires that whatever error the imputation model makes is not itself driven by the treatment $A$ beyond what is captured by $(X, U)$. This is plausible when the imputation model (`Tangram`, `CellPLM`) is fit without access to $A$ (as in our case) and its errors reflect cell-composition or technical effects already summarized by $U$.

Together, `TIDEST`'s claim of valid, orthogonalized inference rests on three conditions: (i) standard double-machine-learning regularity for the cross-fitted nuisance estimators; (ii) $o_p(n^{-1/2})$ consistency of the embedding for the true confounding field; and (iii) asymptotic mean-independence of the imputation-residual correction term from treatment given the estimated confounders.

### B Extended simulation results

#### B.1 Additional data generating processes

Here we provide extended numerical results for the simulation study described in the main text. We compare `TIDEST` with the following methods:

- `DESpace` [2] was run via the `DESpace` Bioconductor package. The function `svg_test()` was called with `verbose = TRUE` to expose the internal `glmLrt` object from which gene-level logFC values were extracted from `glmLrt$table$logFC`. `DESpace` exhibits anti-conservatism at $\alpha = 0$ (FPR$=$ 6.1% vs nominal 5%), consistent with known mild over-dispersion sensitivity of the negative-binomial model on low-count data.

- `SpatialGEE` [3] was run via the `SpatialGEE` R package. $p$-values were obtained from the generalized score test (GST) via `run_gee_gst()`. Effect sizes (Poisson GEE log-coefficients) were obtained from a separate full-model `geeglm()` call using the same $k$-means working correlation structure.

- SpaGCN [4] defines spatial domains and then applies a Wilcoxon rank-sum test between them. We used the signed Wilcoxon $z$-statistic as the effect-size measure.
- For $t$-test, counts were log1p-normalized (no library-size scaling) before the Welch two-sample test. Effect size was the log1p mean difference (superficial minus deep, or treatment minus control).

In addition to the two-region DGP shown in Fig. 2, we evaluated two alternative treatment-generating mechanisms to test robustness of the results:

- **Gaussian random field (GRF) variant.** The treatment $A_i$ is binarized from an independent Gaussian random field with length-scale $\ell = 0.3$, so that the spatial boundary between the two treatment zones is irregular rather than a straight line. This is a harder setting for deconfounding because the treatment and confounder fields have a similar spatial scale. At $\alpha = 1.0$, TIDEST FPR = 7.5% versus SpaGCN 30.7%, $t$-test 33.3%, DESpace 35.4%, and SpatialGEE 27.1%.
- **Latent-factor variant.** The treatment $A_i$ is the binarized first spatial PC (the dominant eigenvector of the Gaussian kernel matrix). This is the most favorable geometry for TIDEST: the treatment is strongly aligned with the leading eigenvector, which is screened out by the correlation threshold $|r| > 0.5$, removing effectively all residual confounding. At $\alpha = 1.0$, TIDEST FPR = 4.0% (near nominal); the $t$-test and SpaGCN also show low FPR (1.7% and 1.4%) in this setting because the spatial structure of the treatment is so dominant that any method with minimal deconfounding benefits. The AUC advantage of TIDEST is largest here (AUC = 0.999 vs 0.984 for SpatialGEE), confirming that TIDEST reconstructs the treatment effect more accurately even when confounding is geometrically easy to remove.

Supp. Fig. B.1 gives FPR, TPR, and AUC at all five confounder levels with standard errors and across DGP variants.

#### B.2 Sensitivity analysis to hyperparameters

##### B.2.1 Augmentation

The augmentation step can be read as a control-variate correction in the spirit of prediction-powered inference [5, 6]. For a target gene $j$, the raw imputation $\tilde{Y}_{ij}$ is a "prediction" whose error $\tilde{Y}_{ij} - Y_{ij}$ is unknown at spot $i$ but is correlated, through the shared imputation model and shared cell-composition structure, with the errors $\tilde{Y}_{ik} - Y_{ik}$ of other genes $k$. For genes $k \in \mathbb{N}_j$ that are *directly observed* in the ST panel, the realized error $Y_{ik} - \tilde{Y}_{ik}$ is known at every spot $i$ and can be used as a control variate: Equation (1) adds back $\tilde{\sigma}_{jk}(Y_{ik} - \tilde{Y}_{ik})$, an estimate of the part of gene $j$'s imputation error explained by gene $k$'s (observed) imputation error, with the external scRNA-seq correlation $\tilde{\sigma}_{jk}$ playing the role of the control-variate coefficient.

Equation (1) averages $|\mathbb{N}_j|$ control-variate terms, each weighted by its own correlation $\tilde{\sigma}_{jk}$ and normalized by $|\mathbb{N}_j|$ (or, with a correlation floor, by the number of neighbors retained). Two competing effects govern the choice of $|\mathbb{N}_j|$ (a parameter whose default is 5): (i) each additional neighbor contributes an independent realization of correlated reconstruction error, so averaging over more neighbors reduces the variance of the correction term, provided the added neighbors remain informative; but (ii) neighbors are ranked by $|\tilde{\sigma}_{jk}|$, so each additional neighbor is more weakly correlated than the last, and once $|\tilde{\sigma}_{jk}|$ for the marginal neighbor is small, including it adds noise from that neighbor's own imputation error without adding much correlated signal. The reconstruction RMSE sweep in Supp. Fig. B.2 (column 1, bottom row) shows exactly this tradeoff empirically: RMSE drops sharply from `top_k` = 1 (0.682) to `top_k` = 5 (0.328, the default), continues to improve modestly to `top_k` = 10 (0.301), and then degrades slightly at `top_k` = 20 (0.326) as increasingly weakly-correlated neighbors are included. TIDEST downstream TPR/FPR/AUC are comparatively insensitive to this choice (TPR $\approx$ 0.92, FPR $\approx$ 0.07 for `top_k` $\in \{5, 10, 20\}$), with the exception of `top_k` = 1, where the much noisier correction reduces power (TPR 0.86).

##### B.2.2 Other hyperparameters

We also vary one parameter at a time around TIDEST default configuration (`top_k` = 5, no correlation floor, no reference noise, 20 SpatialPCA components, PC-treatment screen $|\rho| > 0.5$) – see Supp. Fig. B.2:

- **Correlation floor (`min_corr`).** This parameter drops candidate neighbors with $|\tilde{\sigma}_{jk}|$ below a floor before selecting the top-$k$, normalizing by the number of neighbors actually retained. In the synthetic gene-module

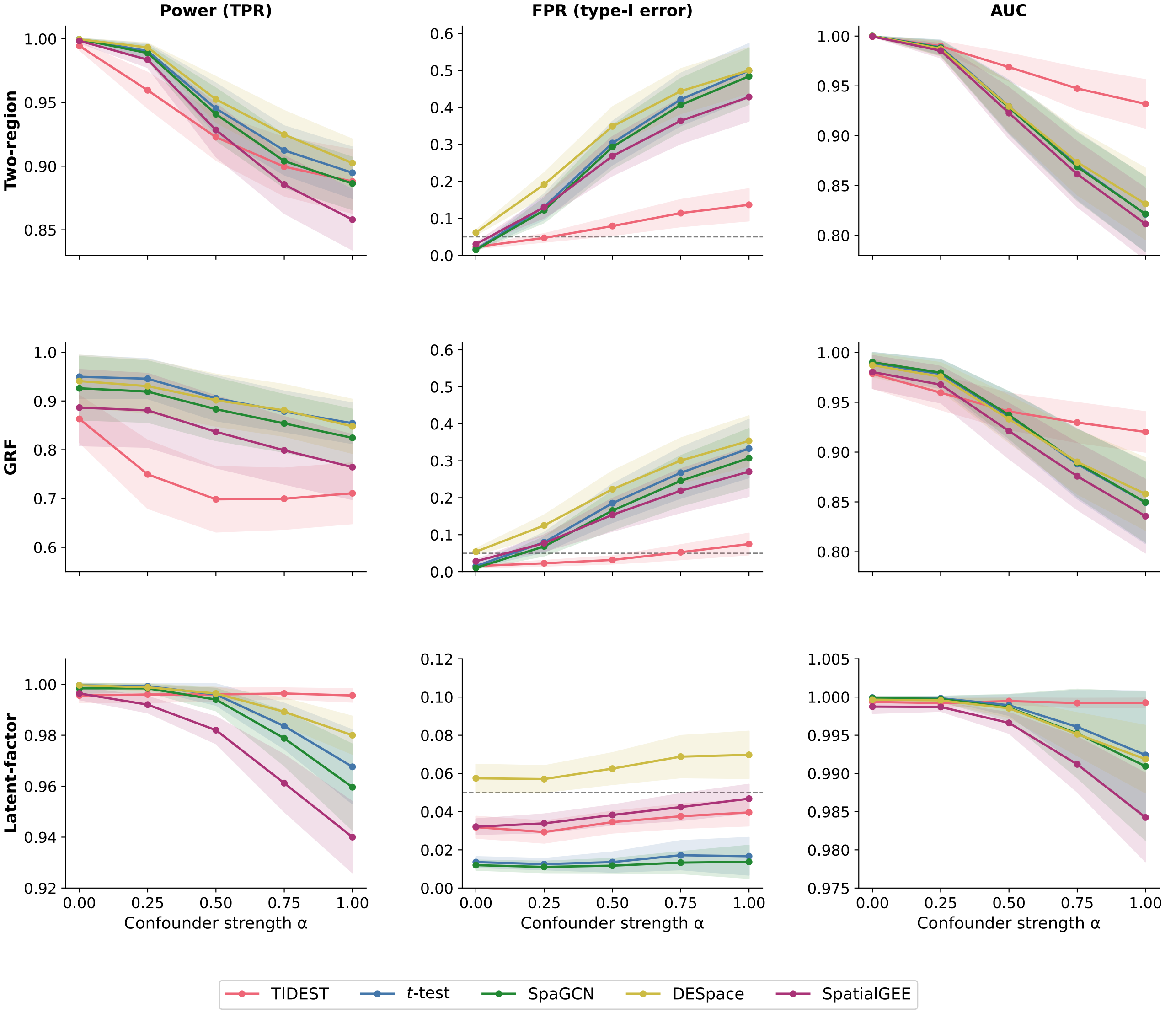


**Figure B.1:** Full simulation results: FPR, TPR, and AUC across all five confounder levels and three DGPs ($\tau_{\text{level}} = 1.0$, $\sigma_{\text{imp}} = 0.5$, $n = 300$ spots, 50 replicates per $\alpha$ level). Shaded bands are 95% confidence intervals across replicates.

structure, the top-5 within-module correlations have a minimum of $\approx 0.41$, so floors up to 0.3 have no effect (identical to the default, RMSE $= 0.328$). Above this range the floor begins to bind: at `min_corr` $= 0.45$, RMSE rises modestly to 0.349; at 0.65, to 0.538 (TPR drops from 0.918 to 0.887). This confirms that a correlation floor is only consequential once it exceeds the typical neighbor correlation, and that `TIDEST` default of using all top-$k$ candidates without a floor is not a meaningfully sub-optimal choice in this regime.

- **Noise on the reference correlation matrix.** We perturbed the scRNA-seq Pearson correlation matrix with symmetric Gaussian noise (SD $\in \{0, 0.05, 0.1, 0.2, 0.3\}$, clipped to $[-1, 1]$) before computing the augmentation, simulating a reference whose gene-gene correlation structure is imperfectly estimated (e.g., from a small or noisy scRNA-seq dataset). Reconstruction RMSE degrades gracefully and roughly monotonically (from 0.328 at SD$= 0$ to 0.435 at SD$= 0.3$), and downstream TPR declines only slightly ($0.918 \to 0.907$) with FPR essentially unchanged ($0.072 \to 0.074$). `TIDEST` reliance on the external reference correlations is therefore robust to substantial mis-estimation of those correlations.
- **Number of `SpatialPCA` components ($n_{\text{pcs}} \in \{5, 10, 20, 50\}$, default 20).** TPR is stable (0.90-0.93) across this range, but FPR is sensitive at the low end: $n_{\text{pcs}} = 5$ gives FPR $= 0.171$, more than double the default's 0.072,

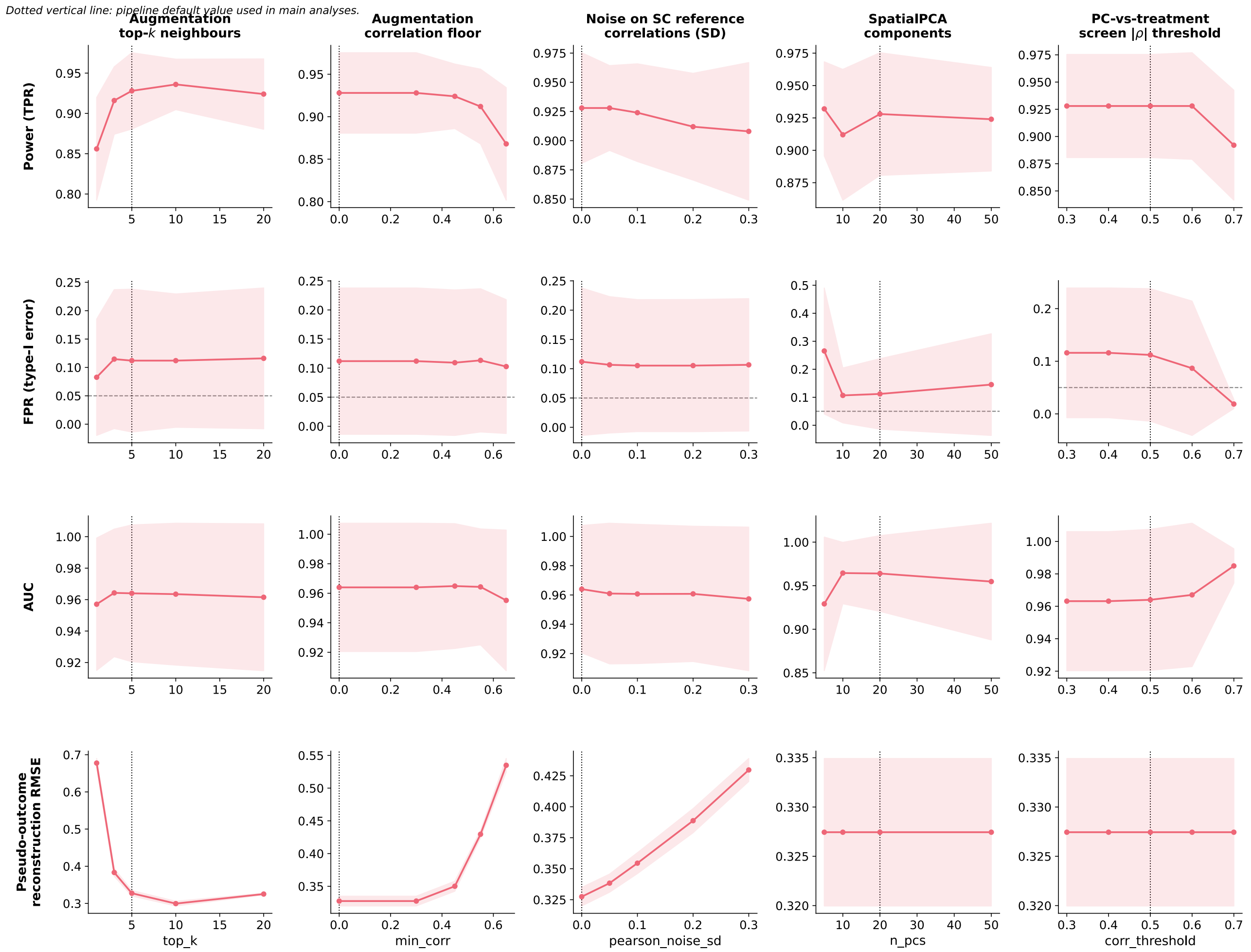


**Figure B.2:** Sensitivity of TIDEST to augmentation and pipeline hyperparameters, at the setting with two-region DGP, $\alpha = 0.5$, $\sigma_{\text{imp}} = 0.5$, $n = 300$, 100 replicates per value. Shaded bands are 95% confidence intervals across replicates. Columns: augmentation neighborhood size (`top_k`), augmentation correlation floor (`min_corr`), additive noise (SD) on the scRNA-seq reference correlation matrix, number of SpatialPCA components, and the PC-vs-treatment correlation screen threshold. Rows: power (TPR), FPR (with the nominal 5% level marked), AUC, and reconstruction RMSE of the augmented outcome $\hat{Y}$ against the true latent expression. Dotted vertical lines mark the default value used in the main analyses.

indicating that too few components fail to capture the spatial confounding field $U$. From $n_{\text{pcs}} = 10$ upward FPR is stable (0.061-0.093), with no further benefit from increasing beyond 20-50.

- **PC-vs-treatment correlation screen ($|\rho|$ threshold $\in \{0.3, \ldots, 0.7\}$, default 0.5).** This threshold controls how aggressively SpatialPCA components correlated with treatment are excluded from the confounder set $\hat{U}$. Lower thresholds (more aggressive exclusion, 0.3-0.4) leave TPR and FPR essentially at the default (0.92-0.93 and 0.075-0.077). Higher thresholds (less aggressive exclusion, retaining components more strongly correlated with treatment in $\hat{U}$) make the procedure more conservative: at 0.7, TPR falls to 0.893 and FPR to 0.050 – closer to nominal but at a power cost. The default 0.5 lies in the flat part of this tradeoff.

### C Mouse brain application details

Interneuron markers serve as negative controls: these cell types are distributed across cortical layers and should not exhibit systematic deep vs. superficial differences. The following table shows BH-adjusted $q$-values for all five interneuron markers across all five methods. TIDEST returns all five as non-significant; *Vip* is called significant by all four competitors.

**Table C.1:** BH $q$-values for interneuron marker genes across five methods. Values in bold are significant ($q < 0.05$).

| Gene | TIDEST | $t$-test | SpaGCN | DESpace | SpatialGEE |
|---|---|---|---|---|---|
| *Pvalb* | 0.096 | 0.355 | 0.482 | 0.618 | 0.469 |
| *Sst* | 0.764 | **0.041** | 0.168 | 0.281 | 0.077 |
| *Vip* | 0.091 | **0.002** | **0.029** | **0.006** | **0.002** |
| *Reln* | 0.140 | 0.125 | 0.259 | 0.260 | 0.154 |
| *Lhx6* | 0.707 | 0.813 | 0.939 | 0.829 | 0.837 |

TIDEST also recovers two genes whose TIDEST estimates align with more recent transcriptomic evidence over older functional annotations. *Pou3f3* (*Brain-1*/*Brn-1*) was initially characterized as a broadly expressed cortical transcription factor [7], but later work established its role in specifying superficial (L2/3) neuron identity [8]: TIDEST detects superficial enrichment ($\hat{\tau} = +0.052$, $q = 3.6 \times 10^{-7}$) that all four competitors miss ($q \geq 0.06$). *Prss12* (*neurotrypsin*), annotated as upper-layer based on biochemical enrichment in dissected superficial fractions [9], is detected by TIDEST as deeply enriched ($\hat{\tau} = -0.066$, $q = 1.1 \times 10^{-10}$), consistent with more recent findings [10, 11]; all competitors agree on direction but only TIDEST reaches significance. In addition to *Fezf2*, the strongest deep-layer signal was *Pcp4* ($\hat{\tau} = -0.198$, $q = 5.4 \times 10^{-69}$), which has well-established deep cortical expression [12].

### D Human glioblastoma application details

Table D.2 lists all 90 genes in the GBM analysis panel, their biological group, expected IvyGAP direction (CT-enriched, LE-enriched, or no clear prior), and the random-effects (DerSimonian-Laird) meta-analysis result across 26 samples. Asterisks indicate BH $q < 0.05$.

**Table D.2:** GBM gene panel with random-effects meta-analysis results ($n = 26$ samples). LE = expected LE-enriched ($\hat{\tau} > 0$); CT = expected CT-enriched ($\hat{\tau} < 0$); 0 = no clear prior. * = BH $q < 0.05$ (random-effects meta-analysis). ✓ = result matches expectation; × = result discordant with expectation; (NS) = not significant ($q \geq 0.05$); – = no directional prior. [a] SALL1 and HEXB are homeostatic microglial markers expected to be LE-enriched. Their CT-enrichment likely reflects a reference artefact: the scRNA-seq reference [13] “Immune” class is overwhelmingly composed of tumor-associated macrophages (TAMs) from the CT compartment, not homeostatic microglia. Tangram therefore maps these cells to CT spots, causing apparent CT-enrichment of homeostatic microglial markers.

| **Gene** | **Exp.** | $\hat{\tau}_{\text{pool}}$ | $z_{\text{pool}}$ | **Result** |
|---|---|---|---|---|
| SNAP25 | LE | +0.247 | +7.3* | ✓ |
| SYT1 | LE | +0.160 | +8.8* | ✓ |
| RBFOX3 | LE | +0.018 | +7.3* | ✓ |
| TUBB3 | LE | +0.055 | +6.6* | ✓ |
| DCX | LE | +0.023 | +6.0* | ✓ |
| DLG4 | LE | +0.006 | +4.1* | ✓ |
| MAP2 | LE | +0.000 | 0.0 | (NS) |
| NRXN1 | LE | +0.023 | +1.8 | (NS) |
| CKB | LE | −0.020 | −1.2 | (NS) |
| MBP | LE | −0.071 | −1.9 | (NS) |
| PLP1 | LE | −0.085 | −2.1 | (NS) |
| MAG | LE | −0.029 | −1.7 | (NS) |
| MOG | LE | −0.045 | −3.0* | × |
| SPARCL1 | LE | +0.067 | +2.1 | (NS) |
| SLC1A2 | LE | +0.077 | +2.3* | ✓ |
| AQP4 | CT | −0.151 | −10.2* | ✓ |
| S100B | CT | −0.128 | −9.1* | ✓ |
| GFAP | CT | −0.147 | −8.3* | ✓ |

| Gene | Exp. | $\hat{\tau}_{pool}$ | $z_{pool}$ | Result |
|---|---|---|---|---|
| APOE | CT | −0.111 | −5.8* | ✓ |
| GJA1 | CT | −0.071 | −5.6* | ✓ |
| GLUL | CT | −0.001 | −0.1 | (NS) |
| ALDOC | CT | −0.009 | −0.3 | (NS) |
| CD44 | CT | −0.092 | −4.0* | ✓ |
| VIM | CT | −0.070 | −3.1* | ✓ |
| VCAN | CT | −0.069 | −5.0* | ✓ |
| CHI3L1 | CT | −0.043 | −1.3 | (NS) |
| FN1 | CT | −0.036 | −1.8 | (NS) |
| TNC | CT | −0.027 | −1.8 | (NS) |
| COL1A1 | CT | −0.007 | −1.8 | (NS) |
| SERPINE1 | CT | −0.003 | −0.3 | (NS) |
| TGFBI | CT | +0.010 | +0.8 | (NS) |
| ANXA2 | CT | −0.048 | −2.0 | (NS) |
| POSTN | CT | −0.003 | −0.6 | (NS) |
| EGFR | CT | −0.108 | −7.4* | ✓ |
| MET | 0 | +0.002 | +0.8 | – |
| ASCL1 | LE | −0.111 | −8.1* | × |
| OLIG1 | LE | −0.126 | −6.9* | × |
| OLIG2 | LE | −0.099 | −6.8* | × |
| PDGFRA | LE | −0.067 | −5.8* | × |
| SOX10 | LE | −0.080 | −5.3* | × |
| SOX4 | LE | −0.043 | −4.7* | × |
| CCND2 | LE | −0.051 | −4.1* | × |
| SOX11 | LE | −0.019 | −3.3* | × |
| EGR1 | LE | −0.025 | −1.8 | (NS) |
| NES | 0 | −0.067 | −6.3* | – |
| BCAN | LE | −0.195 | −7.6* | × |
| PTPRZ1 | LE | −0.119 | −6.3* | × |
| HTRA1 | LE | −0.059 | −5.4* | × |
| NRCAM | LE | −0.048 | −4.5* | × |
| L1CAM | LE | +0.040 | +7.0* | ✓ |
| MMP2 | LE | −0.018 | −4.3* | × |
| ITGB1 | LE | −0.022 | −2.7* | × |
| PLAUR | LE | −0.013 | −0.9 | (NS) |
| CXCR4 | LE | −0.003 | −0.3 | (NS) |
| MMP9 | LE | −0.000 | −0.1 | (NS) |
| MCM6 | CT | −0.023 | −5.2* | ✓ |
| MCM2 | CT | −0.022 | −4.9* | ✓ |
| PCNA | CT | −0.029 | −4.4* | ✓ |
| TOP2A | CT | −0.025 | −3.2* | ✓ |
| MKI67 | CT | −0.017 | −2.4* | ✓ |
| HIF1A | CT | −0.039 | −3.5* | ✓ |
| EPAS1 | CT | −0.016 | −2.6* | ✓ |
| CA9 | CT | −0.005 | −1.0 | (NS) |
| LDHA | CT | +0.007 | +0.5 | (NS) |
| LDHB | LE | −0.061 | −5.6* | × |
| HK2 | CT | +0.000 | 0.0 | (NS) |
| VEGFA | 0 | +0.009 | +1.0 | – |
| CDKN2A | 0 | −0.039 | −3.1* | – |
| IDH1 | 0 | −0.058 | −6.6* | – |
| SALL1 | LE | −0.033 | −6.7* | ×[a] |
| HEXB | LE | −0.015 | −2.4* | ×[a] |

| Gene | Exp. | $\hat{\tau}_{\text{pool}}$ | $z_{\text{pool}}$ | Result |
|---|---|---|---|---|
| OLFML3 | LE | −0.021 | −1.8 | (NS) |
| CX3CR1 | LE | −0.012 | −1.4 | (NS) |
| TMEM119 | LE | −0.007 | −1.1 | (NS) |
| P2RY12 | LE | +0.005 | +1.1 | (NS) |
| SPP1 | CT | −0.059 | −1.8 | (NS) |
| LGALS1 | CT | −0.039 | −1.9 | (NS) |
| LGALS3 | CT | −0.022 | −1.6 | (NS) |
| CD163 | CT | −0.020 | −1.5 | (NS) |
| MSR1 | CT | −0.013 | −1.0 | (NS) |
| MRC1 | CT | −0.002 | −1.3 | (NS) |
| AIF1 | LE | −0.013 | −0.7 | (NS) |
| CD68 | LE | −0.017 | −0.9 | (NS) |
| IL6 | 0 | −0.009 | −3.9* | – |
| PECAM1 | LE | −0.007 | −2.2* | × |
| VWF | LE | −0.004 | −1.1 | (NS) |
| CLDN5 | LE | −0.008 | −1.7 | (NS) |
| PTEN | 0 | +0.010 | +5.0* | – |
| STAT3 | 0 | −0.006 | −1.8 | – |
| PTPN11 | 0 | −0.002 | −0.8 | – |

**Table D.3:** Enrichment direction and $q$-value for the seven OPC/NPC and ECM "paradox" genes, across TIDEST and four competing methods. ✓ indicates the call is significant (BH $q < 0.05$); all reported directions are CT-enriched, the unanimous result for every method–gene combination.

| Gene | TIDEST | $t$-test | SpaGCN | DESpace | SpatialGEE |
|---|---|---|---|---|---|
| *ASCL1* | CT ($8.9 \times 10^{-15}$)✓ | CT (0.003)✓ | CT (0.144) | CT (0.081) | CT (0.065) |
| *OLIG1* | CT ($4.4 \times 10^{-11}$)✓ | CT ($9.7 \times 10^{-7}$)✓ | CT ($3.2 \times 10^{-5}$)✓ | CT ($2.4 \times 10^{-4}$)✓ | CT ($3.6 \times 10^{-7}$)✓ |
| *OLIG2* | CT ($8.2 \times 10^{-11}$)✓ | CT ($1.8 \times 10^{-11}$)✓ | CT ($3.7 \times 10^{-5}$)✓ | CT ($2.1 \times 10^{-5}$)✓ | CT ($4.2 \times 10^{-8}$)✓ |
| *SOX10* | CT ($5.0 \times 10^{-7}$)✓ | CT (0.974) | CT (0.144) | CT (0.426) | CT (0.224) |
| *PDGFRA* | CT ($2.6 \times 10^{-8}$)✓ | CT (0.021)✓ | CT (0.230) | CT (0.412) | CT (0.729) |
| *BCAN* | CT ($3.9 \times 10^{-13}$)✓ | CT ($7.8 \times 10^{-8}$)✓ | CT ($1.2 \times 10^{-9}$)✓ | CT ($5.0 \times 10^{-7}$)✓ | CT ($2.9 \times 10^{-7}$)✓ |
| *PTPRZ1* | CT ($1.6 \times 10^{-9}$)✓ | CT ($4.6 \times 10^{-7}$)✓ | CT ($8.9 \times 10^{-6}$)✓ | CT (0.002)✓ | CT (0.008)✓ |

Beyond *SYT1* and *SNAP25*, several additional normal-brain markers were significantly LE-enriched (Table D.2): *TUBB3* ($q = 3.1 \times 10^{-10}$, significant in 69% of samples), *RBFOX3* (NeuN; $q = 2.5 \times 10^{-12}$, 54%), and *DCX* ($q = 1.1 \times 10^{-8}$, 54%). *SLC1A2* (EAAT2; $\hat{\tau} = +0.077$, $q = 0.041$) was likewise LE-enriched, confirming preserved normal astrocytes at the invasion margin. Regarding the OPC/NPC and ECM "paradox" genes (Fig. **5f**): at the 55 $\mu$m Visium resolution, each CT spot captures tens to hundreds of densely packed GBM tumor cells co-expressing OPC-like markers and ECM proteins, whereas LE spots contain a minority of tumor cells diluted within mostly normal neurons and glia, so the bulk signal is CT-enriched even for genes marking invasive or OPC-like states. This is a property of the data and resolution rather than an artefact of TIDEST deconfounding.

#### D.1 Cross-sample reproducibility as further validation

The biological validations in the main text are evaluated against curated expected directions drawn from the literature. To provide a validation that does not rely on any such prior annotation, we use the 26-sample GBM cohort to ask a purely internal reproducibility question: for each of the 90 panel genes, how often does the sign of the per-sample TIDEST effect estimate $\hat{\tau}$ agree with the sign of the cross-sample, random-effects pooled estimate $\hat{\tau}_{\text{pool}}$? Because the 26 sections come from three independent patient cohorts (MGH, UKF, ZH) with substantial between-tumor heterogeneity (median $I^2 = 93\%$), high sign agreement across samples is not guaranteed by construction and provides

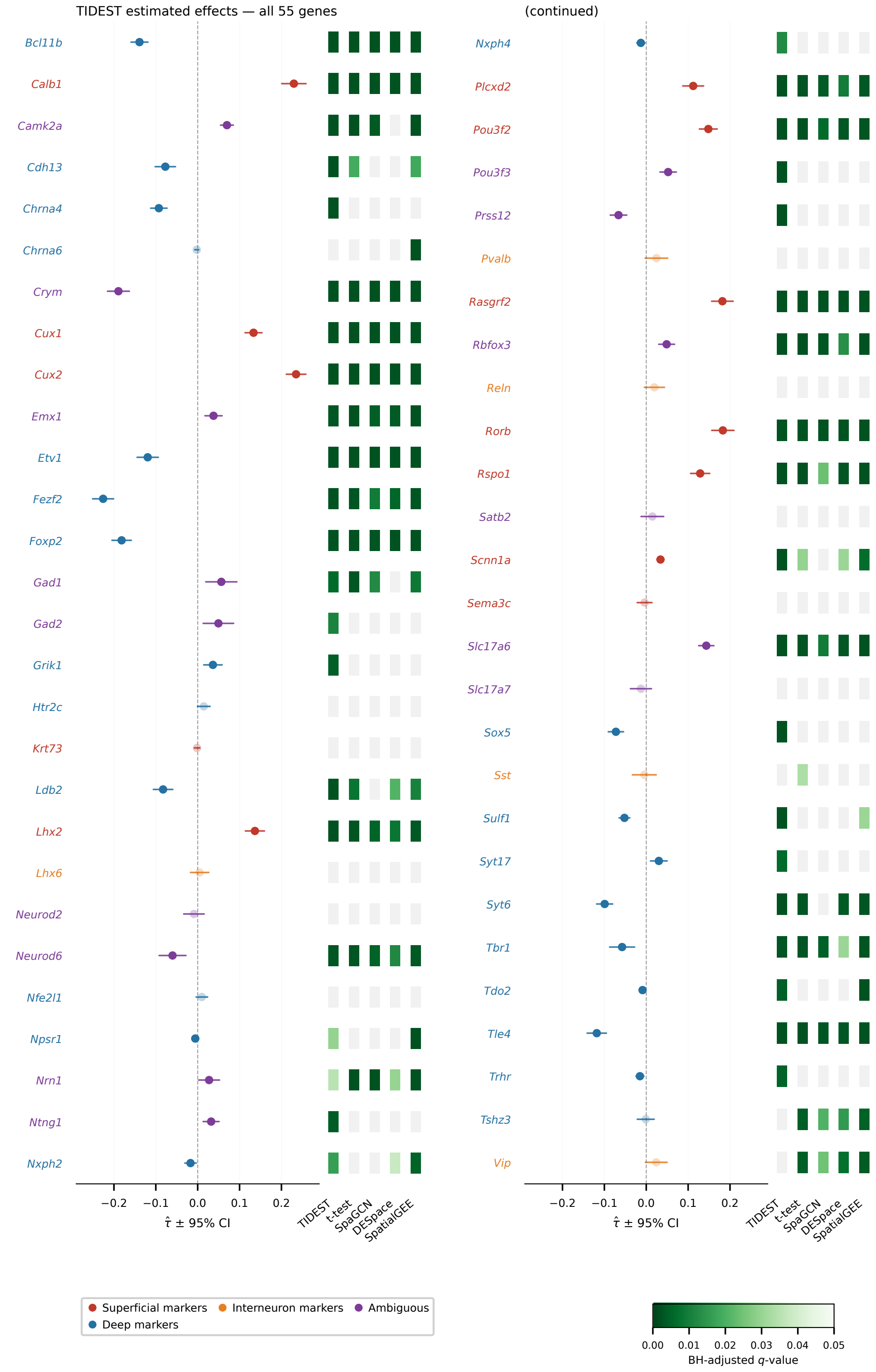


**Figure C.3:** Spatial augmented outcome maps and method comparison for the full mouse brain marker gene panel (55 genes), complementing Fig. **3c-d**.

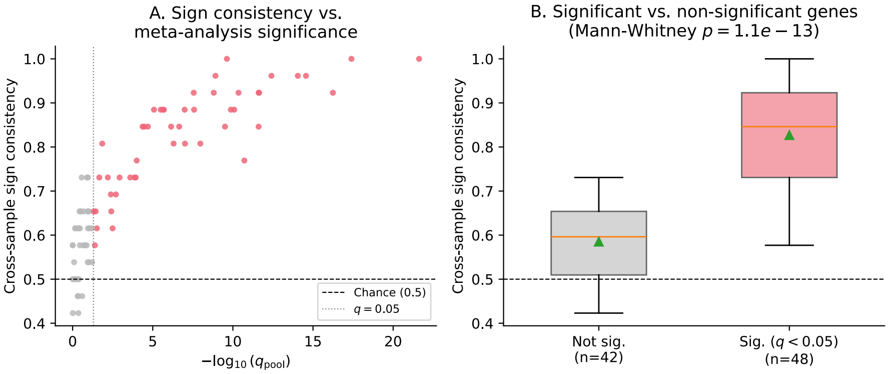


**Figure D.4:** Cross-sample reproducibility of TIDEST effect estimates across the 26-sample GBM cohort, evaluated without reference to curated marker directions. **(a)** Per-gene sign consistency – the fraction of the 26 per-sample $\hat{\tau}$ estimates sharing the sign of the random-effects pooled estimate $\hat{\tau}_{pool}$ – versus meta-analysis significance $-\log_{10}(q_{pool})$. Red: genes significant at $q < 0.05$ ($n = 48$); gray: non-significant ($n = 42$). Dashed line: chance level (0.5); dotted line: $q = 0.05$. **(b)** Distribution of sign consistency for non-significant versus significant genes (boxes show quartiles, orange line the median, green triangle the mean; whiskers extend to the data range). Genes called significant by TIDEST cross-sample meta-analysis show substantially higher cross-sample sign agreement (mean 82.7%) than non-significant genes (mean 58.5%, near chance; Mann-Whitney $p = 1.1 \times 10^{-13}$).

an objective, falsifiable check on whether TIDEST's effect estimates reflect a reproducible biological signal rather than sample-specific noise.

Supp. Fig. D.4 shows, for each gene, the fraction of the 26 samples whose individual $\hat{\tau}$ has the same sign as $\hat{\tau}_{pool}$ ("sign consistency"), plotted against the meta-analysis significance $q_{pool}$ (panel A) and grouped by significance at $q < 0.05$ (panel B). For the 48/90 genes significant in the random-effects meta-analysis, mean sign consistency is 82.7%, compared to 58.5% for the remaining 42 genes – close to the 50% expected by chance for a sign with no reproducible direction (Mann-Whitney $p = 1.1 \times 10^{-13}$). This separation is not an artifact of the significance threshold itself: $q_{pool}$ is computed from a meta-analysis of all 26 samples jointly, whereas sign consistency is a simple per-sample count that could in principle be uncorrelated with the pooled $q$-value if individual estimates were dominated by sample-specific confounding rather than a shared effect. The strong association between the two indicates that TIDEST extracts a direction-consistent signal across independent tumor sections, complementing (and not contingent on) the curated marker-direction comparisons reported in the main text.

### E Human breast cancer application details

**Table E.4:** Per-gene results for the 30 genes present in both the Xenium panel and the curated marker list. **Bold** $\hat{\tau}$ indicates significance ($q < 0.05$) in TIDEST. *Exp. dir.*: expected direction (IBC = IBC-enriched; DCIS = DCIS-enriched; — = contested or no established direction). *Correct*: ✓= significant and correct direction; × = significant but wrong direction; ns = non-significant for a directional gene; — = no established direction. $q$-values are BH-adjusted; values below $10^{-300}$ shown as $< 10^{-300}$.

| Gene | Exp. dir. | $\hat{\tau}$ | $q$ (TIDEST) | $q$ ($t$-test) | $q$ (SpaGCN) | $q$ (DESpace) | $q$ (SpatialGEE) | Correct |
|---|---|---|---|---|---|---|---|---|
| *KRT14* | DCIS | **-0.811** | $< 10^{-300}$ | $< 10^{-300}$ | $< 10^{-300}$ | $< 10^{-300}$ | $<10^{-12}$ | ✓ |
| *ESR1* | DCIS | **-0.669** | $< 10^{-300}$ | $< 10^{-300}$ | $< 10^{-300}$ | $< 10^{-300}$ | $<10^{-12}$ | ✓ |
| *KRT5* | DCIS | **-0.606** | $< 10^{-300}$ | $< 10^{-300}$ | $<10^{-206}$ | $< 10^{-300}$ | $<10^{-13}$ | ✓ |
| *MYLK* | DCIS | **-0.311** | $< 10^{-300}$ | $< 10^{-300}$ | $< 10^{-300}$ | $< 10^{-300}$ | $<10^{-12}$ | ✓ |
| *CDH1* | DCIS | **-0.178** | $< 10^{-300}$ | $<10^{-89}$ | $<10^{-82}$ | $<10^{-62}$ | 0.001 | ✓ |
| *CLDN4* | DCIS | **-0.169** | $<10^{-114}$ | $< 10^{-300}$ | $< 10^{-300}$ | $< 10^{-300}$ | $<10^{-13}$ | ✓ |
| *EGFR* | — | **-0.100** | $< 10^{-300}$ | $<10^{-80}$ | $<10^{-22}$ | $<10^{-77}$ | $<10^{-8}$ | — |

*Continued on next page*

*Table E.4 continued*

| Gene | Exp. dir. | $\hat{\tau}$ | $q$ (TIDEST) | $q$ ($t$-test) | $q$ (SpaGCN) | $q$ (DESpace) | $q$ (SpatialGEE) | Correct |
|---|---|---|---|---|---|---|---|---|
| *PGR* | DCIS | **-0.080** | $< 10^{-300}$ | $<10^{-82}$ | $<10^{-5}$ | $<10^{-128}$ | $<10^{-9}$ | ✓ |
| *GNLY* | IBC | **-0.075** | $<10^{-169}$ | 0.025 | 0.907 | 0.016 | 0.088 | × |
| *S100A8* | IBC | **-0.045** | $<10^{-149}$ | $<10^{-47}$ | 0.003 | $<10^{-70}$ | $<10^{-10}$ | × |
| *CTLA4* | — | **-0.045** | $<10^{-132}$ | 0.055 | 0.907 | 0.008 | 0.225 | — |
| *RUNX1* | DCIS | **-0.038** | $<10^{-20}$ | $<10^{-88}$ | $<10^{-68}$ | $<10^{-83}$ | 0.001 | ✓ |
| *KRT8* | DCIS | **-0.017** | $<10^{-8}$ | $<10^{-286}$ | $<10^{-281}$ | $< 10^{-300}$ | $<10^{-6}$ | ✓ |
| *SNAI1* | IBC | +0.001 | 0.145 | 0.634 | 0.907 | 0.587 | 0.516 | ns |
| *TIGIT* | DCIS | +0.003 | 0.176 | 0.435 | 0.907 | 0.973 | 0.332 | ns |
| *TYROBP* | IBC | **+0.034** | $<10^{-30}$ | $<10^{-7}$ | 0.384 | $<10^{-9}$ | 0.008 | ✓ |
| *KIT* | DCIS | **+0.034** | $<10^{-40}$ | $<10^{-55}$ | $<10^{-5}$ | $<10^{-60}$ | $<10^{-9}$ | × |
| *ZEB1* | IBC | **+0.040** | $<10^{-52}$ | $<10^{-117}$ | $<10^{-15}$ | $<10^{-119}$ | $<10^{-9}$ | ✓ |
| *ZEB2* | IBC | **+0.067** | $<10^{-139}$ | $<10^{-69}$ | $<10^{-17}$ | $<10^{-85}$ | $<10^{-7}$ | ✓ |
| *S100A4* | IBC | **+0.071** | $<10^{-213}$ | $<10^{-22}$ | $<10^{-7}$ | $<10^{-10}$ | 0.003 | ✓ |
| *GATA3* | — | **+0.085** | $<10^{-140}$ | $<10^{-60}$ | $<10^{-133}$ | $<10^{-267}$ | 0.001 | — |
| *NKG7* | IBC | **+0.185** | $< 10^{-300}$ | 0.819 | 0.975 | 0.479 | 0.546 | ✓ |
| *MMP2* | IBC | **+0.207** | $< 10^{-300}$ | $<10^{-160}$ | $<10^{-34}$ | $<10^{-183}$ | $<10^{-8}$ | ✓ |
| *CXCR4* | IBC | **+0.225** | $< 10^{-300}$ | $< 10^{-300}$ | $< 10^{-300}$ | $< 10^{-300}$ | $<10^{-16}$ | ✓ |
| *MLPH* | — | **+0.241** | $< 10^{-300}$ | 0.006 | 0.005 | $<10^{-85}$ | 0.426 | — |
| *ERBB2* | IBC | **+0.268** | $< 10^{-300}$ | 0.169 | 0.907 | $<10^{-138}$ | 0.813 | ✓ |
| *POSTN* | IBC | **+0.279** | $< 10^{-300}$ | $< 10^{-300}$ | $<10^{-234}$ | $< 10^{-300}$ | $<10^{-9}$ | ✓ |
| *FOXA1* | — | **+0.284** | $< 10^{-300}$ | $< 10^{-300}$ | $< 10^{-300}$ | $< 10^{-300}$ | $<10^{-9}$ | — |
| *TOP2A* | IBC | **+0.372** | $<10^{-257}$ | $<10^{-238}$ | $<10^{-190}$ | $<10^{-251}$ | $<10^{-8}$ | ✓ |
| *MKI67* | IBC | **+0.460** | $< 10^{-300}$ | $< 10^{-300}$ | $<10^{-184}$ | $< 10^{-300}$ | $<10^{-11}$ | ✓ |

**Table E.5:** Per-gene results for the 38 genes accessible only through CellPLM prediction (not present in the Xenium 313-gene panel; no competitor benchmarks available). **Bold** $\hat{\tau}$ indicates significance ($q < 0.05$). *Exp. dir.*: as in Table E.4. *Correct*: as in Table E.4; note that with $n = 62755$ cells, even small effects are highly significant – $\hat{\tau}$ magnitude is the primary interpretive metric.

| Gene | Exp. dir. | $\hat{\tau}$ | $q$ (TIDEST) | Correct |
|---|---|---|---|---|
| *TFF3* | DCIS | **-1.082** | $< 10^{-300}$ | ✓ |
| *TFF1* | DCIS | **-0.995** | $< 10^{-300}$ | ✓ |
| *AREG* | DCIS | **-0.734** | $< 10^{-300}$ | ✓ |
| *KRT19* | DCIS | **-0.723** | $< 10^{-300}$ | ✓ |
| *AGR2* | DCIS | **-0.489** | $< 10^{-300}$ | ✓ |
| *CDH3* | DCIS | **-0.281** | $< 10^{-300}$ | ✓ |
| *TP63* | DCIS | **-0.254** | $< 10^{-300}$ | ✓ |
| *LAMB3* | DCIS | **-0.239** | $< 10^{-300}$ | ✓ |
| *LAMC2* | DCIS | **-0.218** | $< 10^{-300}$ | ✓ |
| *ERBB4* | DCIS | **-0.189** | $< 10^{-300}$ | ✓ |
| *CNN1* | DCIS | **-0.186** | $< 10^{-300}$ | ✓ |
| *CLDN3* | DCIS | **-0.134** | $< 10^{-300}$ | ✓ |
| *CLDN7* | DCIS | **-0.116** | $< 10^{-300}$ | ✓ |
| *ITGB6* | DCIS | **-0.053** | $<10^{-174}$ | ✓ |
| *S100A9* | IBC | **-0.041** | $< 10^{-300}$ | × |
| *CA9* | IBC | **-0.005** | $<10^{-78}$ | × |
| *CXCL10* | — | +0.001 | 0.317 | — |
| *ALDH1A1* | IBC | **+0.015** | $<10^{-62}$ | ✓ |
| *COL4A1* | DCIS | **+0.016** | $<10^{-22}$ | × |

*Continued on next page*

*Table E.5 continued*

| Gene | Exp. dir. | $\hat{\tau}$ | $q$ (TIDEST) | Correct |
|---|---|---|---|---|
| *CDKN2A* | IBC | **+0.034** | $< 10^{-300}$ | ✓ |
| *CD44* | IBC | **+0.054** | $< 10^{-300}$ | ✓ |
| *COL11A1* | IBC | **+0.063** | $< 10^{-300}$ | ✓ |
| *MMP9* | IBC | **+0.064** | $< 10^{-300}$ | ✓ |
| *MCM6* | IBC | **+0.078** | $< 10^{-300}$ | ✓ |
| *PCNA* | IBC | **+0.082** | $< 10^{-300}$ | ✓ |
| *GREM1* | IBC | **+0.093** | $< 10^{-300}$ | ✓ |
| *MCM2* | IBC | **+0.102** | $< 10^{-300}$ | ✓ |
| *SNAI2* | IBC | **+0.103** | $< 10^{-300}$ | ✓ |
| *VIM* | IBC | **+0.108** | $< 10^{-300}$ | ✓ |
| *EZH2* | IBC | **+0.109** | $< 10^{-300}$ | ✓ |
| *TWIST1* | IBC | **+0.109** | $< 10^{-300}$ | ✓ |
| *MMP11* | IBC | **+0.126** | $< 10^{-300}$ | ✓ |
| *SPDEF* | DCIS | **+0.201** | $< 10^{-300}$ | × |
| *CDH2* | IBC | **+0.221** | $< 10^{-300}$ | ✓ |
| *THBS2* | IBC | **+0.242** | $< 10^{-300}$ | ✓ |
| *FN1* | IBC | **+0.249** | $< 10^{-300}$ | ✓ |
| *CDK1* | IBC | **+0.259** | $< 10^{-300}$ | ✓ |
| *CCNB1* | IBC | **+0.263** | $< 10^{-300}$ | ✓ |

In addition to *KRT5* and *KRT14*, the myoepithelial program is corroborated by *TP63* ($\hat{\tau} = -0.25$) [14, 15], *MYLK* ($\hat{\tau} = -0.31$) [16], and *CNN1* ($\hat{\tau} = -0.19$) [17], all DCIS-enriched (Tables E.4-E.5). The non-significant immune genes are also biologically interpretable: *TIGIT*, an immune checkpoint receptor [18], shows $\hat{\tau} \approx 0$ ($q = 0.18$), consistent with equivalent immune regulatory pressure in both compartments, and *CXCL10* ($q = 0.32$) shows spatial distribution that does not differ systematically between IBC and DCIS, suggesting locally heterogeneous cytokine signaling that does not resolve into a stable contrast [19]. *NKG7*, a cytotoxic NK/T-cell granule marker [20] expected to be IBC-enriched given immune infiltration is a hallmark of disease progression [21], is detected by `TIDEST` ($\hat{\tau} = +0.185$, $q \approx 0$) but missed by both the $t$-test ($q = 0.77$) and `SpaGCN` ($q = 0.97$).

### References


[1] Jin-Hong Du, Kathryn Roeder, and Larry Wasserman. Assumption-lean post-integrated inference with surrogate-control outcomes. *Biometrika*, page asag004, 2026.

[2] Peiying Cai, Mark D Robinson, and Simone Tiberi. Despace: spatially variable gene detection via differential expression testing of spatial clusters. *Bioinformatics*, 40(2):btae027, 2024.

[3] Yishan Wang, Chenxuan Zang, Ziyi Li, Charles C Guo, Dejian Lai, and Peng Wei. A comparative study of statistical methods for identifying differentially expressed genes in spatial transcriptomics. *bioRxiv*, 2025.

[4] Jian Hu, Xiangjie Li, Kyle Coleman, Amelia Schroeder, Nan Ma, David J Irwin, Edward B Lee, Russell T Shinohara, and Mingyao Li. Spagcn: Integrating gene expression, spatial location and histology to identify spatial domains and spatially variable genes by graph convolutional network. *Nature methods*, 18(11):1342–1351, 2021.

[5] Anastasios N Angelopoulos, Stephen Bates, Clara Fannjiang, Michael I Jordan, and Tijana Zrnic. Prediction-powered inference. *Science*, 382(6671):669–674, 2023.

[6] Ergan Shang, Yuting Wei, and Kathryn Roeder. Predicting the unseen: a diffusion-based debiasing framework for transcriptional response prediction at single-cell resolution. *Proceedings of the National Academy of Sciences*, 122(52):e2525268122, 2025.

[7] Yoshinobu Sugitani, Shigeyasu Nakai, Osamu Minowa, Miyuki Nishi, Kou-ichi Jishage, Hitoshi Kawano, Kensaku Mori, Masaharu Ogawa, and Tetsuo Noda. Brn-1 and brn-2 share crucial roles in the production and positioning of mouse neocortical neurons. *Genes & development*, 16(14):1760–1765, 2002.

[8] Martin H Dominguez, Albert E Ayoub, and Pasko Rakic. Pou-iii transcription factors (brn1, brn2, and oct6) influence neurogenesis, molecular identity, and migratory destination of upper-layer cells of the cerebral cortex. *Cerebral cortex*, 23(11):2632–2643, 2013.

[9] Thomas P Gschwend, Stefan R Krueger, Serguei V Kozlov, David P Wolfer, and Peter Sonderegger. Neurotrypsin, a novel multidomain serine protease expressed in the nervous system. *Molecular and Cellular Neuroscience*, 9(3): 207–219, 1997.

[10] Kazumasa Matsumoto-Miyai, Ewa Sokolowska, Andreas Zurlinden, Christine E Gee, Daniel Lüscher, Stefan Hettwer, Jens Wölfel, Ana Paula Ladner, Jeanne Ster, Urs Gerber, et al. Coincident pre-and postsynaptic activation induces dendritic filopodia via neurotrypsin-dependent agrin cleavage. *Cell*, 136(6):1161–1171, 2009.

[11] Maura Ferrer-Ferrer, Shaobo Jia, Rahul Kaushik, Jenny Schneeberg, Izabela Figiel, Stepan Aleshin, Andrey Mironov, Motahareh Safari, Renato Frischknecht, Jakub Wlodarczyk, et al. Mice deficient in synaptic protease neurotrypsin show impaired spaced long-term potentiation and blunted learning-induced modulation of dendritic spines. *Cellular and Molecular Life Sciences*, 80(4):82, 2023.

[12] Maria Renelt, Viola von Bohlen und Halbach, and Oliver von Bohlen und Halbach. Distribution of pcp4 protein in the forebrain of adult mice. *Acta histochemica*, 116(6):1056–1061, 2014.

[13] Spyros Darmanis, Steven A Sloan, Derek Croote, Marco Mignardi, Sophia Chernikova, Peyman Samghababi, Ye Zhang, Norma Neff, Mark Kowarsky, Christine Caneda, et al. Single-cell rna-seq analysis of infiltrating neoplastic cells at the migrating front of human glioblastoma. *Cell reports*, 21(5):1399–1410, 2017.

[14] Sangjun Lee, Sheila Stewart, Iris Nagtegaal, Jingqin Luo, Yun Wu, Graham Colditz, Dan Medina, and D Craig Allred. Differentially expressed genes regulating the progression of ductal carcinoma in situ to invasive breast cancer. *Cancer research*, 72(17):4574–4586, 2012.

[15] Catalina Lodillinsky, E Infante, A Guichard, R Chaligné, L Fuhrmann, J Cyrta, Marie Irondelle, Emilie Lagoutte, Sophie Vacher, H Bonsang-Kitzis, et al. p63/mt1-mmp axis is required for in situ to invasive transition in basal-like breast cancer. *Oncogene*, 35(3):344–357, 2016.

[16] Dayoung Kim, Jonathan A Cooper, and David M Helfman. Loss of myosin light chain kinase induces the cellular senescence associated secretory phenotype to promote breast epithelial cell migration. *Scientific Reports*, 14(1): 25786, 2024.

[17] Elizabeth Mitchell, Sonali Jindal, Tiffany Chan, Jayasri Narasimhan, Shamilene Sivagnanam, Elliot Gray, Young Hwan Chang, Sheila Weinmann, and Pepper Schedin. Loss of myoepithelial calponin-1 characterizes high-risk ductal carcinoma in situ cases, which are further stratified by t cell composition. *Molecular carcinogenesis*, 59(7):701–712, 2020.

[18] Xin Yu, Kristin Harden, Lino C Gonzalez, Michelle Francesco, Eugene Chiang, Bryan Irving, Irene Tom, Sinisa Ivelja, Canio J Refino, Hilary Clark, et al. The surface protein tigit suppresses t cell activation by promoting the generation of mature immunoregulatory dendritic cells. *Nature immunology*, 10(1):48–57, 2009.

[19] Milim Kim, Hye Yeon Choi, Ji Won Woo, Yul Ri Chung, and So Yeon Park. Role of cxcl10 in the progression of in situ to invasive carcinoma of the breast. *Scientific Reports*, 11(1):18007, 2021.

[20] Susanna S Ng, Fabian De Labastida Rivera, Juming Yan, Dillon Corvino, Indrajit Das, Ping Zhang, Rachel Kuns, Shashi Bhushan Chauhan, Jiajie Hou, Xian-Yang Li, et al. The nk cell granule protein nkg7 regulates cytotoxic granule exocytosis and inflammation. *Nature immunology*, 21(10):1205–1218, 2020.

[21] Sathana Dushyanthen, Paul A Beavis, Peter Savas, Zhi Ling Teo, Chenhao Zhou, Mariam Mansour, Phillip K Darcy, and Sherene Loi. Relevance of tumor-infiltrating lymphocytes in breast cancer. *BMC medicine*, 13(1):202, 2015.
